# Scalable Models of Antibody Evolution and Benchmarking of Clonal Tree Reconstruction Methods

**DOI:** 10.1101/2020.09.17.302505

**Authors:** Chao Zhang, Andrey V. Bzikadze, Yana Safonova, Siavash Mirarab

## Abstract

Affinity maturation (AM) of antibodies through somatic hypermutations (SHMs) enables the immune system to evolve to recognize diverse pathogens. The accumulation of SHMs leads to the formation of clonal trees of antibodies produced by B cells that have evolved from a common naive B cell. Recent advances in high-throughput sequencing have enabled deep scans of antibody repertoires, paving the way for reconstructing clonal trees. However, it is not clear if clonal trees, which capture micro-evolutionary time scales, can be reconstructed using traditional phylogenetic reconstruction methods with adequate accuracy. In fact, several clonal tree reconstruction methods have been developed to fix supposed shortcomings of phylogenetic methods. Nevertheless, no consensus has been reached regarding the relative accuracy of these methods, partially because evaluation is challenging. Benchmarking the performance of existing methods and developing better methods would both benefit from realistic models of clonal tree evolution specifically designed for emulating B cell evolution. In this paper, we propose a model for modeling B cell clonal tree evolution and use this model to benchmark several existing clonal tree reconstruction methods. Our model, designed to be extensible, has several features: by evolving the clonal tree and sequences simultaneously, it allows modelling selective pressure due to changes in affinity binding; it enables scalable simulations of millions of cells; it enables several rounds of infection by an evolving pathogen; and, it models building of memory. In addition, we also suggest a set of metrics for comparing clonal trees and for measuring their properties. Our benchmarking results show that while maximum likelihood phylogenetic reconstruction methods can fail to capture key features of clonal tree expansion if applied naively, a very simple postprocessing of their results, where super short branches are contracted, leads to inferences that are better than alternative methods.

---

Antibodies are Y-shaped proteins consisting of two identical heavy chains and two identical light chains. Antibodies are produced by *B cells* and are used by the immune system to recognize, bind, and neutralize pathogens (also called *antigen*). Unlike other proteins, antibodies are not encoded in the genome directly but present results of somatic *V(D)J recombination* of *immunoglobulin (IG) loci* (Kurosawa and Tonegawa, 1982). Each chain of each antibody is a concatenation of one of V, D (only for heavy chain), and J genes and is konwn as an *IG gene*. An IG gene contains three *complementarity-determining regions* (CDRs) representing antigen binding sites. CDRs are separated by four *framework regions* (FRs) that form a stable structure displaying CDRs on the antibody surface.

After successful binding of an antibody to a given pathogen, the corresponding B cell undergoes the *affinity maturation* (AM) process aiming to improve the *affinity* (i.e., binding ability) of the antibody (Tonegawa, 1983; Neuberger and Milstein, 1995). First, the targeting B cell moves to the *germinal center* (GC) of a lymph node where it undergoes *clonal expansion*: cell divisions that increase the pool of antibodies that bind to the antigen. Then, certain enzymes in the B cell and its clones are activated and introduce *somatic hypermutations* (SHMs) in the utilized IG genes as a means to improve affinity (Muramatsu *et al*., 2000). SHMs change the three-dimensional structure of an antibody (and thus its ability to bind to an antigen) in a stochastic way. The regulatory mechanisms of the immune system play the role of natural selection by expanding B cells with high affinity for antigen and killing self-reactive B cells with potentially harmful mutations. The AM process activates naive B cells (i.e., those that have not been exposed to an antigen) and differentiates them into *memory* and *plasma* B cells. Memory B cells can be repeatedly activated and subjected to the AM, while plasma B cells can secrete massive levels of neutralizing antibodies. Recent studies show that CDRs, which include the binding sites, accumulate more SHMs compared to FRs (Hsiao *et al*., 2019; Safonova and Pevzner, 2019).

The AM process leads to the formation of clonal lineages within a given antibody repertoire, where each clonal lineage is formed by descendants of a single naive B cell. The expressed IG transcripts within the same clonal lineage share a common combination of V, D, and J genes and differ by SHMs only. The evolutionary history of each clonal lineage can be represented by a *clonal tree*, where each vertex corresponds to a B cell and each B cell is connected by a directed edge with all its immediate descendants.

Recent development of sequencing technologies have enabled high-throughput scanning of antibody repertoires (*Rep-Seq*) and have opened up new avenues for studying adaptive immune systems (Georgiou *et al*., 2014; Robinson, 2015; Yaari *et al*., 2015; Watson *et al*., 2017; Miho *et al*., 2018). Rep-Seq technologies enabled AM analysis of antibody repertoires responding to antigens of various diseases: flu (Laserson *et al*., 2014; Horns *et al*., 2019), HIV (Haynes *et al*., 2012; Sok *et al*., 2013a), hepatitis (Galson *et al*., 2016; Eliyahu *et al*., 2018), multiple sclerosis (Stern *et al*., 2014; Lossius *et al*., 2016), rheumatoid arthritis (Elliott *et al*., 2018). Such analysis allows biologists to identify broadly neutralizing antibodies (Yermanos *et al*., 2018) and reveal antigen-specific and general mutation patterns (Horns *et al*., 2019; Hsiao *et al*., 2019).

An intriguing feature of the clonal trees is that due to the short time frame they represent, they can differ from phylogenetic trees. Some of the sequenced nodes may belong to the internal nodes of the tree instead of the tips. Also, there is no reason to assume that the tree should be bifurcating or even close to bifurcating. Thus, unlike traditional phylogenetics, perhaps Steiner trees (which can put observations at *some* of the internal nodes) or spanning trees (that put an observation at *all* internal nodes) should be preferred for reconstructing antibody sequences (Fig. 1a). Various reconstruction methods have been developed attempting to recover clonal trees from antibody sequences (e.g., Jiang *et al*., 2013; Sok *et al*., 2013b; Lee *et al*., 2017; Hoehn *et al*., 2017; Horns *et al*., 2016; Lees and Shepherd, 2015; DeWitt *et al*., 2018). Some of these methods use simple clustering methods (e.g., Jiang *et al*., 2013), while others formulate the problem as a Steiner tree problem (Sok *et al*., 2013b; Lee *et al*., 2017; Horns *et al*., 2016; DeWitt *et al*., 2018) or maximum-likelihood (ML) phylogenetic tree reconstruction under models of sequence evolution (Hoehn *et al*., 2017; Lees and Shepherd, 2015).

**Fig. 1.**
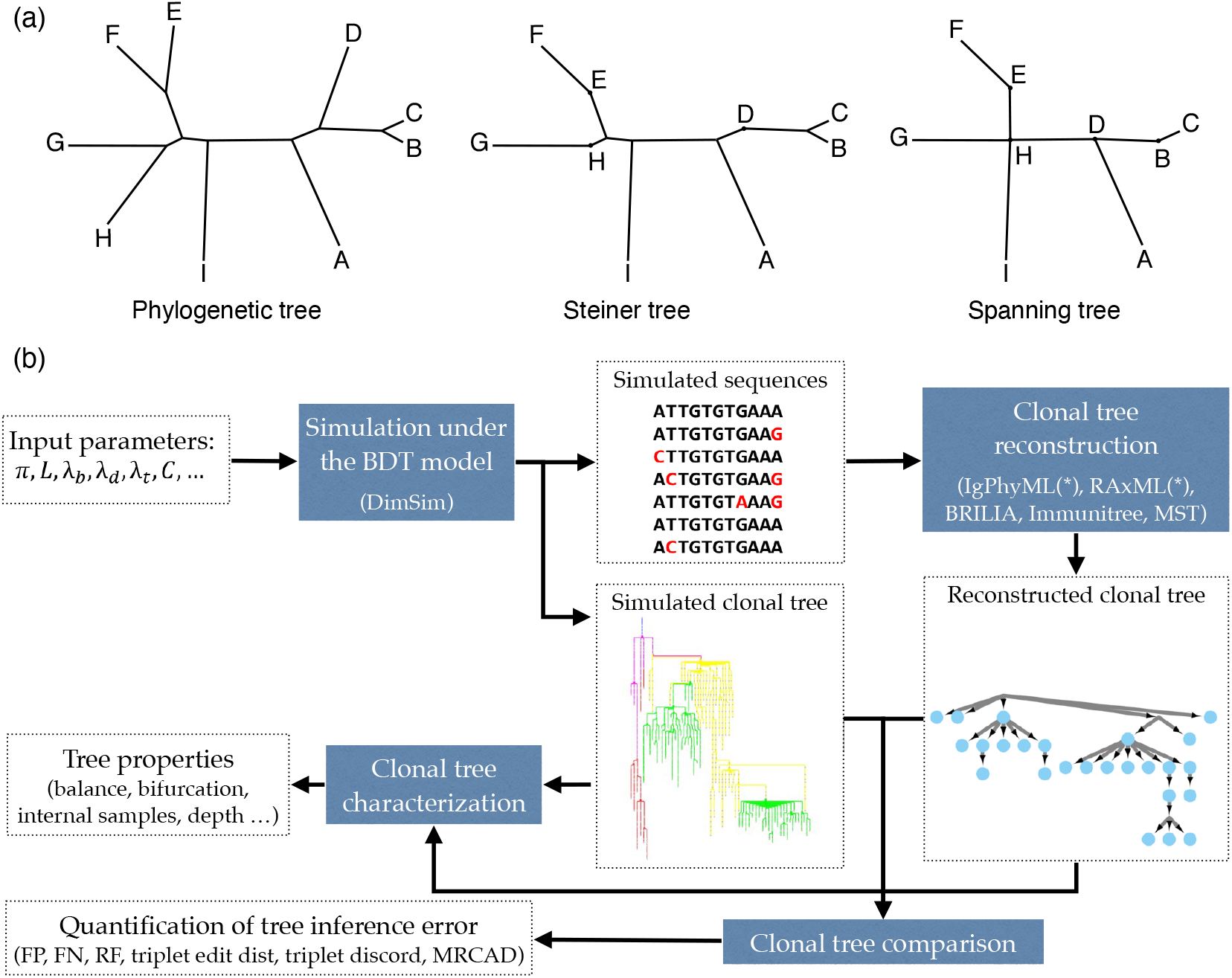
(a) Examples of a phylogenetic tree, a Steiner tree, and a spanning tree. Letters indicate sequenced data. Phylogenetic trees put all data points at leaves and none at internal nodes, spanning trees put data at every node (whether internal or leaf), and Steiner trees are in between (some but not all internal nodes correspond to data). (b) The evaluation framework. The BDT model, parameterized by several values (Table 1) is first sampled using the fast algorithm implemented in DIMSIM to create the simulated (i.e., “true”) sequence data and clonal trees. These trees are then reconstructed from the simulated sequence data using various methods. The reconstructed clonal tree is compared to the simulated tree using several metrics adopted here to account for internal node sampling and multifurcation. Properties of true and inferred trees are measured using metrics such as balance and resolution.

In order to evaluate methods proposed for reconstructing clonal trees, we need models for antibody sequence evolution and clonal tree expansion that can be used for simulation. This modeling step is challenging for several reasons. *i*) Since selection is an important force in AM, it needs to be modelled directly, or else, the shape of the resulting trees will not be realistic. Traditional phylogenetics simulations first simulate a tree of sampled taxa and then evolve sequences down the tree. This two-step approach simplifies simulation but is not sufficient for AM because the strong selection effects make the evolution of the clonal tree and the antibody sequences interdependent. A better approach is to co-evolve the tree and *all* evolving sequences. The challenge in co-evolving is to design a principled model for how sequences impact evolution and to develop a scalable simulation algorithm that can generate millions of cells (which can then be subsampled). *ii*) Literature suggests that there are hotspots and coldspots of SHMs (e.g., Rogozin and Kolchanov, 1992; Pham *et al*., 2003). However, traditional models of sequence evolution are i.i.d and will miss the context-dependence. *iii*) Different types of antibody cells (e.g., activated and memory cells) have very different mutational and selection behaviors and these distinctions need to be modelled.

There have been several attempts at designing models that are appropriate for clonal expansion in AM (e.g., Childs *et al*., 2015; Amitai *et al*., 2017; Reshetova *et al*., 2017; Davidsen and Matsen, 2018). As many processes involved are complex and hard to model exactly, these models have all taken different routes. For example, determining affinities of sequences to hypothetical antigens is difficult, as affinity binding itself is a complicated chemical process, and each method models affinity in a different fashion. Nevertheless, all these methods have limitations, which we will return to in our discussion session. In summary, they do not scale to very large number of cells (millions), they allow for simulating one round of infection (as opposed to an evolving antigen and recurring infections), and they do not model various types of antibody B cells. We propose that to simulate realistic clonal trees and correctly benchmark lineage reconstruction tools, we need models that are generic and flexible, so that they can be updated as a better understanding of the underlying processes is developed. One goal of the present work is to provide such a scalable and flexible simulation framework. In addition to simulation, we note that comparing clonal trees and characterizing their properties require extending metrics from phylogenetics to trees with internal node samples and multifurcations.

In this paper, we make several contributions. *i*) We introduce a general birth, death, transformation (BDT) model and describe how BDT can be instantiated to create a model of AM that simultaneously co-evolves the clonal tree and antibody sequences. *ii*) We introduce a scalable sampling algorithm for our model that enables generating very large trees (millions of cells). *iii*) We refine existing metrics and define new ones for characterizing properties (e.g., balance) of clonal trees and define a set of evaluation metrics for comparing them. *iv*) We study a small post-processing step applied to ML phylogenetic inference and show that it effectively deals with the problem of internal node sampling in antibody sequences. *v*) We perform extensive simulation studies (Fig. 1b) under various parameters and benchmark the performance of seven reconstruction methods: minimum spanning tree, existing tools BRILIA (Lee *et al*., 2017), IgPhyML (Hoehn *et al*., 2017), RAxML (Stamatakis, 2014), and Immunitree (Sok *et al*., 2013b), and modified methods IgPhyML* and RAxML*). We study how the parameters of the AM model impact properties of clonal trees and reconstruction error. These studies showcase the power and flexibility of our benchmarking framework.

## Generative Model

We first define a general Birth/Death/Transformation (BDT) model and introduce an efficient algorithm for sampling trees from the BDT model. We then instantiate the general model for simulating AM processes.

### The birth/death/transformation (BDT) model

Forward-time birth-death models are used extensively in macro-evolutionary modelling (Nee, 2006), whereas micro-evolution simulations often use coalescent models, which hope to approximate forward time evolution, albeit not always successfully (Stadler *et al*., 2015). We start by describing a general forward-time model that can allow realistic micro-evolutionary simulations by ensuring that birth and death rates are not constant, and instead change with properties of evolving units (e.g., cells).

#### Model description

In the BDT model, a set of *particles* continuously undergo birth (B), death (D), and transformation (T) events. Each particle *i* has a list of properties 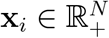. At each moment in time, the system contains a set *S* of *n* active particles, and each active particle *i* ∈ *S* undergoes birth, death, and transformation events according to independent Poisson point processes. In the birth event, a particle *i* is removed from *S* and new particles *j* and *k*, with properties **x**_*j*_ and **x**_*k*_, are added to *S*; properties **x**_*j*_ and **x**_*k*_ are drawn from a distribution determined by **x**_*i*_ and model parameters. In the event of the death for particle *i*, it is removed from *S*. In the transformation event, a particle *i* is removed from *S* and a new particle *j* with properties **x**_*j*_, drawn from a distribution determined by **x**_*i*_, is added to *S*. Starting from a single node and continuously applied, this process defines a rooted tree where nodes are all particles that ever existed (including those that died); birth events create bifurcations, transformation events create nodes with one child, and death events create leaves with no child. The tree can be subsampled as desired.

For each particle *i* ∈ *S*, the birth rate, death rate, and transformation rate are thoroughly determined by its properties **x**_*i*_ and 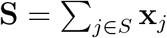, the sum of property vectors over all particles. We let Λ_*B*_(**x**_*i*_, **S**), Λ_*D*_(**x**_*i*_, **S**), and Λ_*T*_ (**x**_*i*_, **S**) denote the birth, death, and transformation rates, respectively. In the time interval between two events for any two particles in the system, we assume a memoryless process. Thus, these rates remain constant between any two events but can change when an event happens. The ratio between the birth rate and the death rate, both of which are functions of the particle properties, can be thought of as the factor controlling the selective pressure, which can be time-variant.

Because of the memoryless property, the time until the next BDT event always follows the exponential distribution with rates Λ_*B*_(**x**_*i*_, **S**), Λ_*D*_(**x**_*i*_, **S**), and Λ_*T*_ (**x**_*i*_, **S**) for each event type. The time until any event for any particle follows an exponential distribution with *λ* =Σ_*i*∈*S*_ (Λ_*B*_(**x**_*i*_, **S**) + Λ_*D*_(**x**_*i*_, **S**) + Λ_*T*_ (**x**_*i*_, **S**)). The probability of the next event being a specific event *E* ∈ {*B, D, T* } for a particular particle *i* is 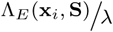. Specifying the rate functions and the distribution of properties at the initial state fully specifies the model.

#### Efficient sampling under the general model

The model we described can be efficiently sampled if we also assume that we are able to write 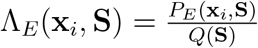 where 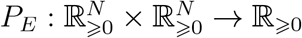 and 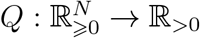 are polynomial functions with a constant degree, where coefficients of *P*_*E*_ are non-negative. Thus, for any particle *i* ∈ *S*, the birth rate can be written as 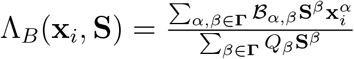 where **Γ** = [0 … *γ*]^*N*^ for some integer *γ*, ℬ_*α,β*_ and *Q*_*β*_ are coefficients of the polynomials, and **a**^**b**^ denotes 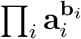 for vectors **a** and **b**. We can write Λ_*D*_(**x**_*i*_, **S**) and Λ_*T*_ (**x**_*i*_, **S**) similarly by replacing ℬ_*α,β*_ with 𝒟_*α,β*_ and 𝒯_*α,β*_.

With this assumption, 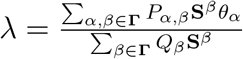 where *P*_*α,β*_ = ℬ_*α,β*_ + 𝒟_*α,β*_ + 𝒯_*α,β*_ and 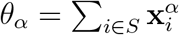 for all *α* values (note that **S** = *θ*_1_). Thus, to efficiently sample the time till the next event, we only need *θ*_*α*_ values which we can simply store and update in constant time after each event. This allows for a constant time sampling of the next event time (in terms of *n*) for constants *N* and *γ*. Once we sample the time till the next event, we need to sample one of the three possible events. The probability of the next event being birth for particle *i* is (derivations shown in the supplementary material)

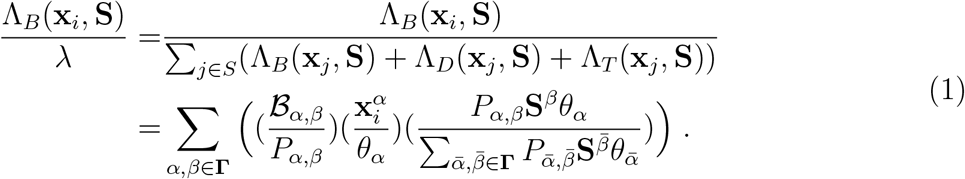

and probability of each death and transformation events can be written similarly.

We now suggest the following sampling procedure (see Algorithm S1):

1. Sample (*α, β*) pair (representing one term of the polynomial) from a multinomial distribution on **Γ** *×* **Γ** where each pair has probability 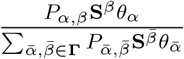.
2. Sample particle *i* from a distribution on *S* where each *i* has probability 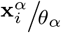.
3. Sample birth, death, or transformation with probabilities 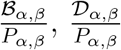, and 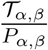.

In this procedure, the probability of selecting the birth event for a particle *i* is simply 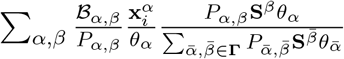, which matches Equation (1) (ditto for death and transformation events). Step 1 takes constant time (in terms of *n*) given that *θ*_*α*_ values (and thus **S**) are pre-computed for all *α*; step 2 can be achieved in *O*(log *n*) time using an interval tree data structure to store partial sums of 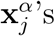 (see Algorithm S1); step 3 takes constant time. Thus, a tree on *k* nodes drawn from the distribution defined by the BDT process can be sampled in *O*(*k* log(*k*)) time by repeated applications of Algorithm S1.

### Antibody Affinity Maturation (AM) model

We now define a specific case of the general model for dynamic antibody affinity maturation. Our goal is to model how antibody-coding sequences evolve in response to several rounds of infections by an evolving antigen (e.g., flu). Simulations according to this AM model are implemented in a C++ tool called Dynamic IMmuno-SIMulator (DIMSIM).

In this paper, we focus on simulating the heavy chain sequences only (thus, by antibody-coding sequences we mean only the heavy chains). While light chains might be important for some immunological applications, most existing Rep-Seq studies focus on sequencing heavy chains only (e.g., Stern *et al*., 2014; Ellebedy *et al*., 2016; Magri *et al*., 2017; Horns *et al*., 2019). Also, since only memory B cells can be repeatedly activated by the encounter with an antigen, we will simulate memory B cells only. Plasma B cells do not undergo SHMs and represent terminal states of the clonal lineage development and thus can be sampled from the leaves of the simulated tree if needed. We will refer to a B cell that has just encountered an antigen and moved to a GC as an *activated B cell* (Fig. 2a).

**Fig. 2.**
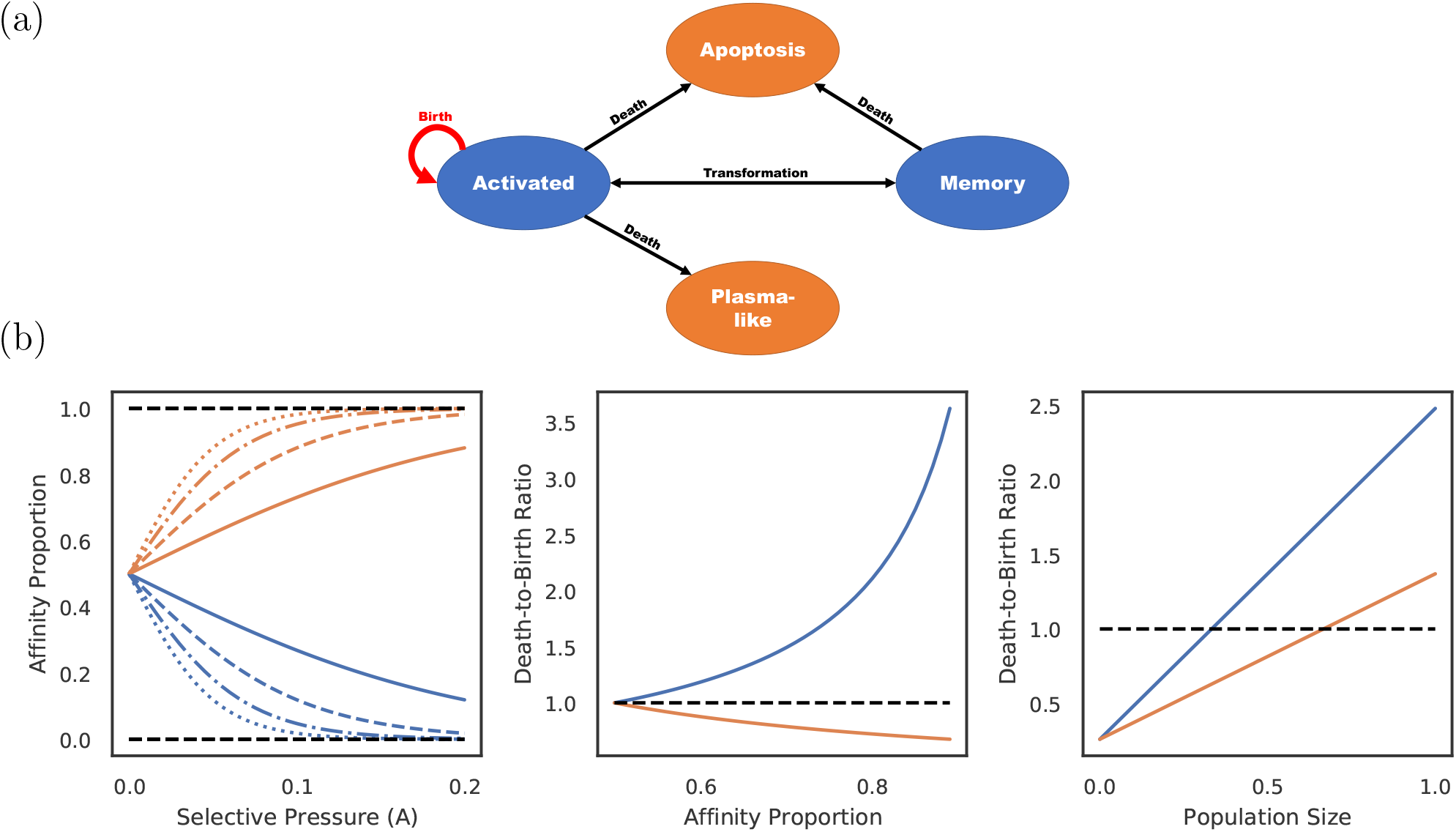
(a) States of cells and transitions during infected stage. Only states colored blue are modeled. Transitions to states colored orange are treated as death events. (b) Consider a population of activated B cells where all cells have one of two sequences: L (Blue) or H (Orange). Let *ρ* be the ratio of affinity of H-type cells to L-type cells, and let the affinity proportion be the total affinity of H (or L) cells over the affinity of all cells (i.e., 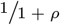 for H and *ρ/*1 + *ρ* for L). Left: The affinity proportion as a function of the selective pressure *A* when the sequence closeness to the target is *f*_*ζ*_(*L*) = −50 and *f*_*ζ*_(*H*) = −10, −20, −30, or −40 (respectively: dotted, dashed/dotted, dashed, or solid). Middle: the ratio of death rate to birth rate as a function of affinity proportion of H cells, fixing the population size to the carrying capacity. Right: ratio of death rate to birth rate as a function of the population size normalized by the carrying capacity, fixing *ρ* = 2. All other parameters set to defaults (Table 1). The selective pressure *A* and the level of binding control the portion of affinity taken up by better sequences (left), which controls the growth of the cell type (middle), but the growth rate is also a function of the total population size (right).

#### Rounds and stages

The model simulates *r* rounds of infection. Each round consists of two stages, an *infected* stage, where a set of new antigens initiate a response that activates the B cells being modeled, and a *dormant* stage, where the B cells being modeled are not actively involved in an immune response. The generative model is identical in the two stages but is parameterized differently. The system can switch between the two stages using user-defined rules including those that reflect infection progression (described below). During the infected stage of round *i*, we assume the existence of a *given* target amino-acid sequence of length 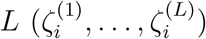 (without any stop codon), defined as the best possible antibody-coding sequence that can bind to the present antigen. When antigens evolve from one round to the next, the target should also change. The model has many parameters related to the immune system properties (Table 1), which we define as we progress.

**Table 1.** Parameters of the AM model

| Parameter name | Default value | Parameter description |
| --- | --- | --- |
| $\lambda'_d$ | $1/402$ | Rate (inverse life time) of cell death for memory cells ($\text{days}^{-1}$) |
| $\lambda_b$ | 6 | Rate of cell division for activated B cells ($\text{days}^{-1}$) |
| $\lambda_d$ | $10^4$ | Rate of cell death during dormant stage ($\text{day}^{-1}$). |
| $\lambda_t$ | 0.01 | Rate of activation of a typical responsive memory cell |
| $\rho_p$ | $1/100$ | Portion of activated B cells that turn into plasma cells per cell division |
| $\rho_m$ | $1/4$ | Portion of activated B cells that turn into memory B cells per cell division |
| $\mu$ | $5 \times 10^{-4}$ | Rate of SHMs per base pair per generation |
| $\mathbf{K}^5$ | See appendix | Empirical 5-mer mutation frequencies per generation |
| $L$ | 125 | Length of the amino acid antibody-coding sequence (assuming the length is fixed) |
| **CDR** | $\{31 \dots 35, 50 \dots 65, 98 \dots 114\}$ | Positions of the three CDR regions (amino acid coordinates) |
| $\delta(i, j)$ | Table S1 | BLOSUM matrix defined on a pair of amino-acids $i$ and $j$ |
| $\Delta_0$ | -120 | BLOSUM score of a typical responsive memory B cell antibody-coding sequence to target |
| $\Delta'_0$ | -75 | BLOSUM score of activated B cell antibody-coding sequences that leads to cure |
| $w_f$ | $1/3$ | BLOSUM score multiplier of non-CDR positions (i.e., FRs) |
| $\kappa$ | 2 | BLOSUM score ratio of antibody-coding sequences to antigen sequences |
| $A$ | 0.1 | Selective pressure: factor connecting sequence similarity and log binding affinity |
| $\rho_a$ | $1/2$ | Factor connecting log affinity and B cell activation (sensitivity to affinity level $A$) |
| $C$ | $10^5$ | Carrying capacity limited by total resources (see text for meaning) |
| $M$ | $Ce^{A\Delta'_0}$ | The threshold of the sum of affinity for a stage change |
| $r$ | 56 | Rounds of viral infections |
| $\hat{\Psi}$ | See appendix | Nucleotide sequence of the initial B cell |
| $\zeta_1, \dots, \zeta_r$ | See appendix | Target amino acid sequences for viral infections in each round |
| $\eta_1, \dots, \eta_r$ | See appendix | Flu sequences assumed as antigens in the simulation |
| $t_1, \dots, t_r$ | See appendix | Starting time of each infected stage (day) |

#### Cell Properties

In the AM model, each particle *i* represents a B cell with the property vector **x**_*i*_ = (*g*_*i*_, *s*_*i*_, *t*_*i*_, *g*_*i*_ / *a*_*i*_, *g*_*i*_*a*_*i*_). The binary property *g*_*i*_ = 1 indicates whether a cell *i* has entered a germinal center of a lymph node, in which case we call it an activated B cell (or “activated cell” for short); *g*_*i*_ = 0 indicates a memory B cell outside lymph nodes, which we call a “memory cell” for simplicity. The *s*_*i*_ property encodes the DNA sequence of B cell *i* coding for the variable region of the heavy chain with a fixed length 3*L* (for the sake of simplicity, we assume the faith of the cell depends only on the variable region of the heavy chain). The other properties are derived from the first two properties, but we keep them as part of **x**_*i*_ because they allow us to define Λ_*E*_(**x**_*i*_, **S**) functions as polynomials of saved properties (Table 2); this, in turns, enables the use of our fast sampling algorithm. Property *t*_*i*_ denotes the rate of transformation. For memory cells, *t*_*i*_ is the rate at which the memory cell activates and becomes an activated cell in response to an antigen. For activated cells, *t*_*i*_ is the rate at which the activated cells mature into memory cells. Thus, transformations, which only happen during the infected stage, create a child cell *j* with property *g*_*j*_ = 1 − *g*_*i*_ and *s*_*j*_ = *s*_*i*_. Property *a*_*i*_ denotes the strength of affinity binding of the Ig receptor of the cell *i* to the antigen. We let 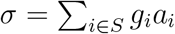 denote the fifth element of vector **S**; thus, *σ* is the total affinity of activated cells and *a*_*i*_/*σ* is the fraction of total affinity assigned to a cell. We will show how *t*_*i*_ and *a*_*i*_ are set based on the sequence of *i* and the target. The fourth and fifth properties are simple functions of other properties.

**Table 2.**
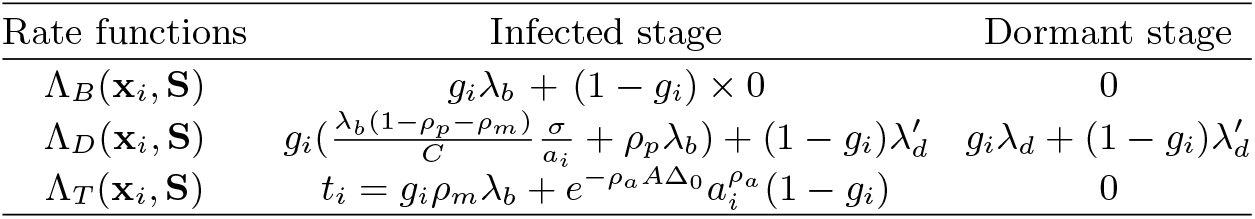
Birth, death, and transformation rates. See Table S3 for polynomial forms.

| Rate functions | Infected stage | Dormant stage |
| --- | --- | --- |
| $\Lambda_B(\mathbf{x}_i, \mathbf{S})$ | $g_i \lambda_b + (1 - g_i) \times 0$ | 0 |
| $\Lambda_D(\mathbf{x}_i, \mathbf{S})$ | $g_i \left( \frac{\lambda_b(1 - \rho_p - \rho_m)}{C} \frac{\sigma}{a_i} + \rho_p \lambda_b \right) + (1 - g_i) \lambda'_d$ | $g_i \lambda_d + (1 - g_i) \lambda'_d$ |
| $\Lambda_T(\mathbf{x}_i, \mathbf{S})$ | $t_i = g_i \rho_m \lambda_b + e^{-\rho_a A \Delta_0} a_i^{\rho_a} (1 - g_i)$ | 0 |

#### Sequence evolution

Each cell has a fixed sequence, and mutations occur at the time of a cell birth, which happens only for activated cells in the infected stage. After a birth event for cell *i*, properties *s*_*j*_ and *s*_*k*_ of child cells *j* and *k* are chosen independently and identically at random. While any sequence evolution model could be incorporated in the DIMSIM framework, we describe below a 5-mer-based model used in these analyses.

#### Determining sequence affinity

Affinity *a*_*i*_ is only defined and used during the infected stage where the target is available (it is undefined during the dormant stage). We define the affinity *a*_*i*_ of a cell *i* as a function of its sequence *s*_*i*_ and the target sequence *ζ*. The closer the sequence to the target, the higher its affinity should be. Exact relationships between the sequences and affinity are not know and cannot be easily modelled. For the purpose of benchmarking, any reasonable function should suffice. Assuming *f*_*ζ*_ (*s*_*i*_) gives a measure of closeness of the sequence to the target in the affinity space, we set

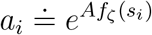

where *A* is a constant factor used to calibrate the selective pressure (see below). We will describe our particular choice of function *f*_*ζ*_ (*s*_*i*_) using BLOSUM similarity below.

#### Rates

The event rates are functions of cell properties and the stage (Table 2). During the dormant stage, there are no births or transformations; cell only die with a very high uniform rate *λ*_*d*_ for activated cells and a low uniform rate 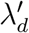 for memory cells.

During the infected stage, we adjust death rates of cells based on their affinities but keep the birth rates constant; this interplay is used to simulate the selective pressure. An activated cell can undergo cell division at a uniform rate *λ*_*b*_, differentiate into a memory cell at a uniform rate *t*_*i*_ = *ρ*_*m*_*λ*_*b*_ or a plasma-like cell at a uniform rate *ρ*_*p*_*λ*_*b*_ driven by helper T cells, and undergo apoptosis (i.e., die) driven by follicular dendritic cells (FDCs). We do not model plasma-like cells; instead, both differentiation into plasma-like cells and apoptosis are treated as death events (Figure 2a). The rate of apoptosis of activated cell *i* is inversely proportional to the amount of resources (antigens and FDCs) to which cell *i* has access when competing against other activated cells. Thus, the proportion of resources available to cell *i* is modelled by the affinity proportion *a*_*i*_/*σ* (i.e., the affinity of the cell to the antigen normalized by the current sum of the affinity of all activated cells). This affinity proportion is impacted by the choice of parameter *A*. The lower the *A*, the more uniform these proportions become, as expected with low selective pressure; conversely, as *A* increases, *a*_*i*_/*σ* values further diverge between low affinity and high affinity cells (Fig. 2b). Thus, *A* can be used to control the strength of the selective pressure.

The memory cells undergo apoptosis at a uniform rate 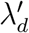. They can also activate by helper T cells to enter the germinal center and become an activated cell at the rate *t*_*i*_ set to:

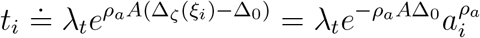

Note that activation rate of memory cells increases monotonically with their affinity to the target, according to 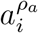 where *ρ*_*a*_, set by default to 1*/*2, is the sensitivity of B cell activation to affinity. This dependency on affinity is to model the increased propensity of the memory cells to activate when presented by helper T cells with familiar antigen. The choice of the default *ρ*_*a*_ = 1*/*2 is motivated by the fact that although memory cells with higher binding strengths to the antigen are more likely to be activated, the interaction between a helper T cell and a memory B cell is an one-time event, and is thus less sensitive to binding strength.

As an example, consider a system with two cell types: L and H, each type with its own unique sequence (Figure 2b). Assume all cells are activated cells, the number of L and H are the same at one point in time, and H cells have a higher affinity than L cells by a factor of *ρ*. For ease of exposition, here, we include mutation rate as part of the death rate because mutation events also decrease cell count. Let’s assume the total number of cells equals the carrying capacity *C*. If L and H have the same affinity (i.e., *ρ* = 1), then the birth and death rates are identical for all cells. As the affinity of H cells is increased (i.e., *ρ >* 1), the death rate of L cells increases linearly whereas the death rate of H cells decreases. Thus, H cells will have higher birth rates than death, will be selected for, and will expand. If we fix *ρ* = 2 and increase the population size, the death rates of both L and H cells increase but at different rates. When the population size is small compared to *C*, both types of cells have more birth than death. After a threshold (*C/*3 in this example), the death rate of L type surpasses its birth rate (thus, its population starts to shrink) while the population of H cells continues to grow. However, as the population size increases (2*C/*3 here), both sets of cells start to shrink (i.e., higher death rates than birth).

### Default Models Choices

Several steps of our simulations are flexible and can be changed by the user to provide reasonable models. We next describe the particular choices we made in our experiments below, noting that these choices can be changed.

#### Stopping criteria

The system enters dormant stage when antigens are neutralized by the antibodies. A simple way to define neutralization is to switch the stage when the total affinity of antibodies produced by plasma-like cells reach a certain threshold; here, we switch when the sum of affinities of activated cells (*σ*) reaches a predefined constant *M*.

#### Sequence evolution

In our experiments, we use an empirical 5-mer-based model inspired by Yaari *et al*. (2013). Let 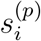 be the nucleotide on the *p*-th position of nucleotide sequence of cell *i*. Each 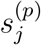 or 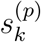 is independently set to *s* ∈ {*A, C, G, T* } with probability:

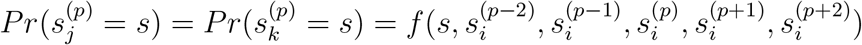

where *f* : *{A, C, G, T }*^6^ → [0, 1] denotes an empirically determined 5-mer frequency model based on the model of Yaari *et al*. (2013) and recomputed based on newer datasets including non-synonymous mutations (see details in the supplementary material).

#### Modelling affinity

While various methods can be imagined for measuring closeness of the sequence to the target, we used a simple approach: measuring sequence similarity according to the BLOSUM matrix and appropriate scaling of numbers. In this formulation, we assume each amino-acid position contributes to the binding strength to the target and the sanity of the structure of Ig-receptor independently. Thus, we model affinity proportionally to the product of the effect of each amino-acid position. This simple model completely ignores the 3D structure of proteins, but we argue, is sufficient for the purpose of creating benchmarking datasets.

When *s*_*i*_ includes a stop codon, we simply set *a*_*i*_ = 0. Otherwise, let 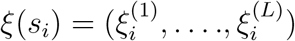 denote the antibody-coding amino-acid sequence of cell *i*. We define the BLOSUM score of an amino acid sequence *ξ* as

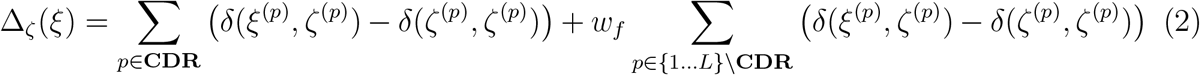

where *δ*(., .) gives the BLOSUM score between two amino acids (Table S1), and *w*_*f*_ is a constant used to calibrate the importance of CDRs versus FRs in the affinity and transformation processes. We then simply set *f*_*ζ*_ (*s*_*i*_) = Δ_*ζ*_ (*ξ*(*s*_*i*_)).

#### Choosing targets

Several rounds of target sequences are assumed to be provided as input, and the extent of the change in targets across rounds impacts the patterns of the immune response and hence the shape of the clonal trees that result. In our experiments, to define targets across rounds, we seek a set of sequences with an evolutionary trajectory that reflects the evolutionary history of a set of real antigen (e.g., influenza virus). Let the known amino-acid sequences of an antigen sampled through time (flu sequence over seasons) be denoted by ***η***_1_, …, ***η***_*r*_, and let each sequence have the fixed length *L*_*η*_. To choose the targets, we first select an arbitrary naive B cell, here chosen from datasets of Ellebedy *et al*. (2016), and set 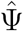 to antibody-coding nucleotide sequence of the variable region of its heavy chain. Then, we simply set ***ζ***_1_ to the amino-acid translation of 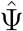. In other words, in the first round, we use the naive cell as the target, and therefore, the first couple of rounds of the simulation should be treated as dummy rounds and should be discarded. Let *κ* be a positive constant that controls the rate of change in the target relative to the rate of change in the antigen sequences. To define the remaining targets, we seek to find the set of *r* − 1 sequences that minimize:

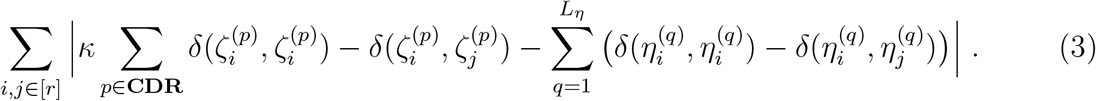

This score simply penalizes a set of targets by the divergence between pairwise sequence distances of all target sequences across all rounds versus pairwise sequence distances of all antigen sequences over the same rounds. To account for the presence of conserved regions, we arbitrarily chose to keep all the non-CDR regions invariable in all target sequences (note that the chose of invariable sites can be easily changed). Thus, if the score is minimized, the distance between two target sequences from two rounds would become similar to the distances of antigen sequences, scaled by a factor of *κ*. We approach this NP-hard problem using a greedy search heuristic (Algorithm S3). The heuristic starts with arbitrary ***ζ***_2_, …, ***ζ***_*r*_ and replaces one symbol of one sequence at a time to reduce the objective function; it repeats until reaching a local minimum where no such replacement is possible.

## Materials and Methods

### Flu simulations

#### Simulation settings

We performed several simulations of a series of *r* = 56 rounds of flu infections, using sequences of hemagglutinin (HA) protein. HA found on the surface of the influenza viruses is the primary target of neutralizing antibodies. High mutation rate of influenza genome changes the sequence of HA and allows the virus to escape from the immune pressure thus making flu recurring seasonal infection. The NCBI Influenza Virus Resource (Bao *et al*., 2008) contains 961 HA sequences from influenza B virus collected around the world. Each HA sequence is labeled with a year and a location. For simulation purposes, we extracted 59 HA sequences corresponding to flu infections in Hong Kong and selected 56 out of 59 HA sequences that have the same length (584 aa). The selected HA sequences were detected in Hong Kong from 1999 to 2010.

We used the default settings for the various parameters of Table 1, and used the approach described earlier to choose the target amino-acid sequences. Each round corresponds to one season, starts at the infected stage with a given target sequence ***ζ***_*l*_, which ends when *σ* = *M*. At that point, we assume the infection is overcome and the system switches to dormant, where we stay until the next round starts (times of flu outbreaks are known in our dataset). When the *r* = 56 rounds of infections end, we sample *ζ* = 200 antibody-coding nucleotide sequences Ψ_1_, …, Ψ_*ζ*_ from cells in the system (i.e., from the round *r*) and built their clonal tree.

#### Experiments

To benchmark reconstruction tools, we set up four experiments, varying one or two parameters in each experiment (Table 3) and setting the remaining ones to default values (Table 1). The central experiment contains 19 conditions, changing the selective pressure (*A*) and the rate of hypermutation (*µ*). We vary *A* from 1*/*8*×* of default value (0.1) to 2*×* and vary *µ* s from 1.25 *×* 10^−4^ to 2 *×* 10^−3^ per base-pair per generation. In six combinations, the selective pressure is not high enough to overcome random mutations; in these cases, the affinity values do not increase and as a result, the carrying capacity is never reached. Thus, we exclude these conditions. We also study three other parameters. We vary the weight multiplier of FRs (*w*_*f*_) from 1*/*5 to 2. We vary the carrying capacity (*C*), which is the germinal center size or the amount of antigens FDCs hold in the context of B cell maturation, from 12500 to 400000. The value of this parameter can impact the speed of novel mutations arising and may change the properties of simulated trees. We also vary the mean life-time of memory cells from 0.5 year to 16 years, to study the impact of the extent of memory cell activation during recurrent infections.

**Table 3.** Experiment setup

| Experiment | Controlled parameters | Parameter values | Parameter units |
| --- | --- | --- | --- |
| Selective pressure vs. rate of hypermutation | $A \times \mu$ | $(2, 2), (2, 1), (2, 1/2), (2, 1/4), (2, 1/8), (1, 2), (1, 1), (1, 1/2), (1, 1/4), (1, 1/8), (1/2, 1), (1/2, 1/2), (1/2, 1/4), (1/2, 1/8), (1/4, 1), (1/4, 1/2), (1/4, 1/4), (1/4, 1/8), (1/8, 1/4), (1/8, 1/8)$ | $A : 10^{-1},$<br>$\mu : 10^{-3}$ |
| Framework weight | $w_f$ | $2, 1, 1/2, 1/3, 1/5$ | 1 |
| Germinal center size | $C$ | $4, 2, 1, 1/2, 1/4, 1/8$ | $10^5$ |
| Memory cell life | $1/\lambda'_d$ | $16, 8, 4, 2, 1, 1/2$ | year (365 days) |

### Methods of Clonal Lineage Reconstruction

#### MST(-like) methods

We implemented a simple minimum spanning tree method in the following way. We let the vertices of the graph to correspond to Ψ_1_, …, Ψ_*ζ*_ as well as 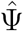. For each pair of vertices, we let the distance between them to be the Hamming distance between corresponding nucleotide distance. We then find the minimum spanning tree (MST) of the graph and root the resulting tree at the vertex corresponding to 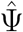.

Besides MST, we also ran reconstruction using Immunitree Sok *et al*. (2013b), a tool that clusters antibody-coding sequences into lineages and builds clonal trees at the same time by optimizing a minimum spanning tree and Steiner tree-like problem. We took as input Ψ_1_, …, Ψ_*ζ*_ and used Immunitree to build a set of clonal trees. We then added vertex 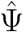 as the root and let the roots of the clonal trees to be immediate children of 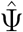.

*Brilia* clusters antibody-coding sequences into lineages and builds clonal trees at the same time. We took as input Ψ_1_, …, Ψ_*ζ*_ and used Brilia to build a set of clonal trees. We then added vertex 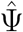 as the root and added roots of the clonal trees as children of 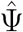.

#### Phylogenetic methods

We tested ML based phylogenetic reconstruction using on RAxML under GTR model and IgPhyML, a ML method tuned specifically for immune cells. For RAxML, we took as input Ψ_1_, …, Ψ_*ζ*_ as well as 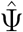 to obtain an unrooted phylogenetic tree and reroot at 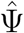. For IgPhyML, we took as input Ψ_1_, …, Ψ_*ζ*_ and provided 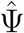 as root to obtain a rooted phylogenetic tree. Both methods produce fully binary trees.

#### Zero-aware phylogenetic methods

Since the length of each antibody-coding nucleotide sequence 3*L <* 400, it is reasonable to assume that both ends of any branch with length less than 10^−4^ would correspond to the same sequence (if it was sampled). Therefore, we slightly modified RAxML and IgPhyML by contracting branches of length less than 10^−4^ and we call the new methods RAxML* and IgPhyML* respectively.

### Evaluation Framework

#### Notations

The simulated and reconstructed histories of samples Ψ_1_, …, Ψ_*ζ*_ are represented as trees where samples are uniquely labeled on some nodes and the remaining nodes are left unlabelled. For a rooted tree *T*, we let **L**_*T*_ be the set of leaves and **I**_*T*_ be the set of internal nodes. For each node *v* of *T*, let *C*(*v*) be the set of its children. We define *ϕ*(*v*) as the set of node labels of labeled nodes below *v*. Also, for any *set* of nodes *V*, we define *ϕ*(*V*) = *{ϕ*(*v*) : *ϕ*(*v*) ≠ ∅, *v* ∈ *V* } and *ϕ*(*T*) = *ϕ*(**I**_*T*_ *∪* **L**_*T*_).

#### Characterizing a clonal tree

We define a set of metrics for characterizing properties of simulated trees in terms of their topology, branch length, and distribution of labelled nodes (Table 4). Some of these metrics are motivated by similar ones on phylogenetic trees but are adjusted to allow sampled internal nodes and multifurcations.

**Table 4.** Properties of a clonal tree T.

| Property | Definition |
| --- | --- |
| Internal sample (%) | The percentage of labeled nodes in set $\mathbf{I}_T$. |
| Bifurcation index | Defined as $\frac{\vert \mathbf{I}_T\vert }{\vert \mathbf{L}_T\vert -1}$ equals 1 for bifurcating trees and $\approx 0$ for the star tree. |
| Sample depth | The average depth of labeled nodes in $T$. |
| Balance (cherry) | Half the sum over all leaves of the fraction of their siblings that are also leaves.<br>$\sum_{v \in \mathbf{I}_T} \binom{\vert \mathcal{C}(v) \cap \mathbf{L}_T\vert }{2} / (\vert \mathcal{C}(v)\vert - 1)$ where $0/0 \doteq 1/2$ |
| Single mutation branches (%) | The percentage of branches with length one. |
| Accumulated mutations (avg) | The average depth (path length to the root) of all labeled nodes of tree $T$. |
| Accumulated mutations (sum) | The summation of branch lengths of all branches of tree $T$. |
| Mutations per branch | The average branch length of tree $T$. |
| The last four metrics require branch length (in mutation unit) on the tree. |  |

For example, to measure tree balance, we extend the definition of the number of cherries but allow modifications (our definition reduces to the traditional definition when the tree is binary). Other metrics (e.g., percent internal samples) are only meaningful for clonal trees and are meant to quantify the deviation of a clonal tree from phylogenetic trees.

#### Comparing trees

Many metrics exist for comparing phylogenetic trees. However, in the presence of polytomies and sampled ancestral nodes, the classic metrics need to be amended. Here, we generalize several existing metrics and introduce new ones. All metrics are defined over a simulated tree *R* and a reconstructed tree *E*, both induced down to include all labeled nodes (i.e., removing unlabelled nodes if less than two of their children have any labelled descendants). See Table 5 for precise definitions of metrics.

**Table 5.**
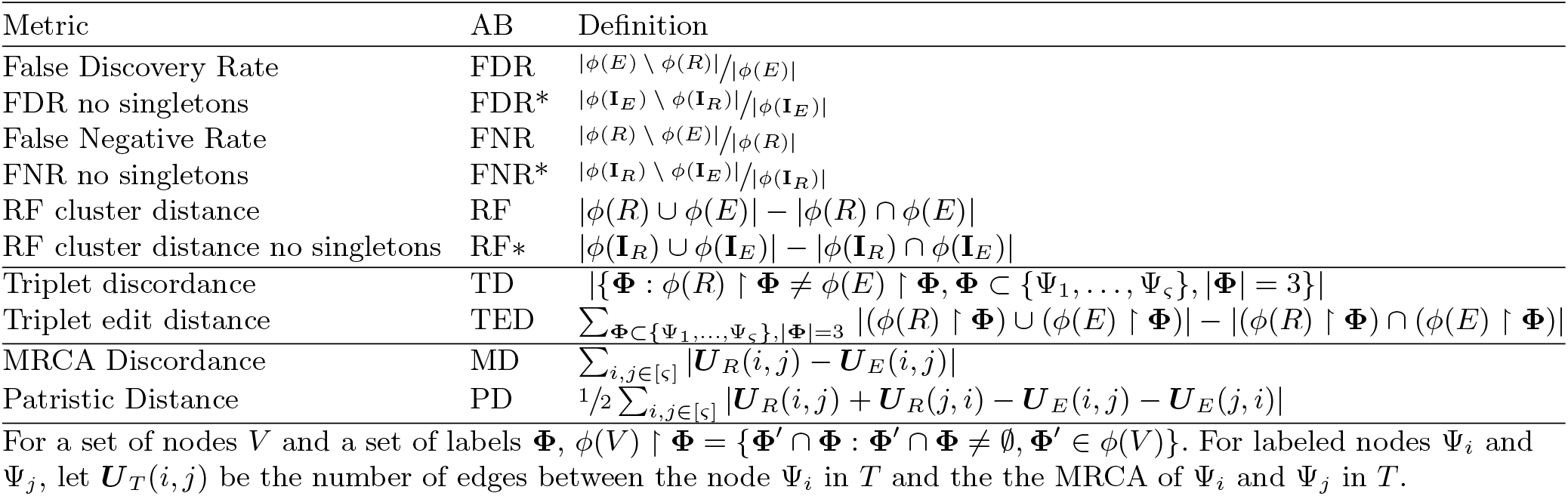
Metrics for comparing the reference simulated tree R to estimated tree E.

#### RF-related

We define False Discovery Rate (FDR) as the percentage of clusters in *E* that are not in *R*, False Negative Rate (FNR) as the percentage of clusters in *R* that are not in *E*, and Robinson-Foulds cluster distance (RF) as the number of clusters in either but not both trees. Note that unlike traditional Robinson and Foulds (1981) distance, here, internal nodes can also have labels, and we define the metric based on clusters in a rooted tree instead of bipartitions in an unrooted tree. Moreover, the singleton clusters are trivial when all labeled nodes are leaves; however, when there are labeled internal nodes, including or excluding singletons can make a difference. Thus, we also define FPR FNR, and RF distance when excluding singleton clusters.

#### Triplet-based

We define *triplet discordance* (TD) as the number of trees induced by triples of *labeled* nodes (leaf or internal) where the topology in the simulated tree and the reconstructed tree differ. We define the *triplet edit distance* (TED) as the summation over all triplets of the labeled nodes of cluster RF distance between the two trees induced to the triplet. Intuitively, it is the sum of the minimum number of branch contractions and resolutions required to covert a triplet in *R* to a triplet in *E*, summed over all triplet.

#### Path discordance

Patristic discordance for a pair of labelled nodes Ψ_*i*_ and Ψ_*j*_ is defined as the difference between the number of branches in the path between Ψ_*i*_ and Ψ_*j*_ on two trees *R* and *E*. The patristic discordance (PD) between *R* and *E* is the summation of the Patristic discordance over all pairs of labelled nodes (intern or leaf). We define the MRCA discordance for an ordered pair of labelled nodes Ψ_*i*_ and Ψ_*j*_ as the difference between the number of branches in the path between Ψ_*i*_ and its MRCA with Ψ_*j*_ when computed from trees *R* and *E*. The MRCA discordance (MD) between the two trees is the summation of MRCA discordance over all ordered pairs of labeled nodes.

The FNR and FDR metrics are already normalized. To normalize other metrics, for each experimental condition, we create a control tree by randomly permuting labels of the true tree. We then normalize scores (other than FNR and FDR) of a reconstruction method by dividing it by the average score of replicates of the control method.

Computing FNR, FDR, and RF metrics takes *O*(*ζ*) time with hashing and randomization (algorithm S4). Triplet-based metric can be easily computed in *O*(*ζ*^3^) time with simple preprocessing and iterating over all triplets. Both PD and MD take *O*(*ζ*^2^) time with preprocessing that computes distances to MRCAs.

## Results

### Demonstration of the simulation process

Visualizing one replicate of simulation under default condition, we see patterns of average affinity and the number of activated and memory cells that rise and fall as time progress during the infected stage (Fig. 3a). During each round of infection, the affinity first decreases and then increases as long as the duration of the infection is long enough. This pattern agrees with biological expectations: when the number of activated cells is low and the selective pressure is low, a mutation is likely to lead to reduced affinity, whereas, when the number of activated cells increases, the selective pressure begins to increase and select for higher affinity. The duration of infections, the mean affinity at the end, and the total number of cells also varies widely across different seasons. When the affinity at the start of a season is low, the duration of infection is longer and more activated cells and memory cells are generated (Figs. 3a and S1a). This pattern is also consistent with the biological expectation: when the immune system already has high affinity to the antigen, it can eradicate the antigen quickly and without much need for further evolution. To further quantify the pattern, we define the novelty of each target *ζ*_*i*_ as the negation of the maximum BLOSUM score between that target and any previous target: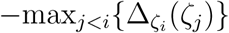. We observe that as novelty of the target increases, the average affinity of activated cells at the end of the infection tends to decrease (*R*^2^ = 0.242, *p* = 2.5 *×* 10^−4^), whereas, the number of activated cells at the end of the infection (*R*^2^ = 0.248, *p* = 2.0 *×* 10^−4^) and the duration of infection (*R*^2^ = 0.288, *p* = 4.8 *×* 10^−5^) both tend to increase (Fig. 3b).

**Fig. 3.**
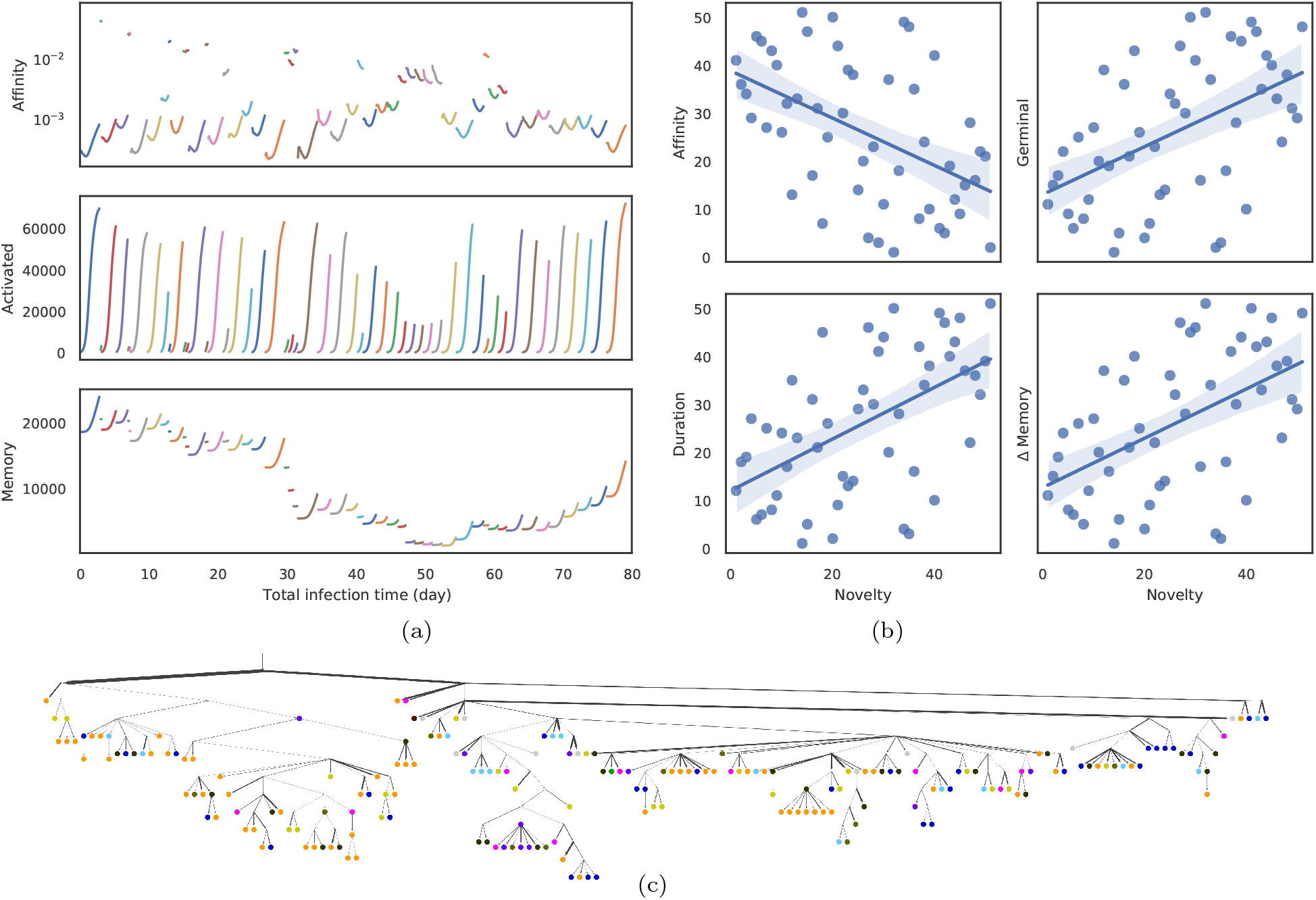
a) Average affinity of activated cells to current infection target (log scale), the number of activated cells, and the number of memory cells by total time in infected stage across the last 51 stages of infection (colors) each corresponding to one flu season (discarding the first 5 rounds and dormant stages). b) Impact of the novelty of the antigen on the outcome of the infection across the 56 seasons simulated. The novelty of seasons is measured by 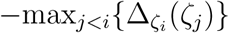 and is ranked from less novel to more novel on the x axis. Y-axis shows ranking (from low to high) of average affinity of activated cells to current infection target (*R*^2^ = 0.242, *p* = 2.5 *×* 10^−4^) at the end of the infection, the number of activated cells (*R*^2^ = 0.248, *p* = 2.0 *×* 10^−4^) at the end of the infection, the duration of infection (*R*^2^ = 0.288, *p* = 4.8 *×* 10^−5^), and the change in memory cell count (*R*^2^ = 0.264, *p* = 1.2 *×* 10^−4^) from the start to the end of the infection. c) Clonal tree of memory cells sampled from one simulation under default condition after all 56 seasons. Nodes are colored by seasons when the memory cells emerge (grey for season 1 through 46; as part (a) for others). Here, 17 internal nodes are sampled and are indicated as circles. Edge weights denote the number of mutations of sequences denoted by adjacent nodes. See Figure S1 for more.

Memory cells counts fluctuate. Each season leads to a buildup in memory cells from the start to the end of the infection, and the amount of buildup depends on the duration and correlates with novelty (*R*^2^ = 0.264, *p* = 1.2 *×* 10^−4^). However, the total number of memory cells reduces between seasons due to cell deaths (Fig. S1c) and changes across seasons. In particular, a string of short-lived infections and large time spans between the flu seasons between 2002 and 2008 gradually lead to a depletion of the memory cells, which are then built up again in the subsequent seasons (Fig. S1c).

### Benchmarking reconstruction methods

#### Default Parameters

Under default parameters, over all evaluation metrics, zero-aware Phylogenetic methods (IgPhyML* and RAxML*) clearly have the best accuracy in reconstructing the lineage history (Fig. 4). The normal Phylogenetic methods (IgPhyML and RAxML), which produce fully binary trees with no samples at leaves, have the lowest FNR error, retrieving more than 90% of the correct clusters. However, their precision is predictably low: close to 35% of their clusters are incorrect. Interestingly, zero-aware phylogenetic methods have only a slight increase in FN rate (*<* 2% on average) but enjoy a dramatic improvement in precision. By simply contracting super-short branches, the FDR error reduces to less than 15%, which is better than all other methods. Similarly, normal phylogenetic methods perform poorly according to RF, PD, and MD metrics, which emphasize false positives, but perform well (but not as well as the zero-aware versions) according to triplet-based metrics (TED and TD), which penalize false negatives more than false positives. Among the two phylogenetic reconstruction methods, RAxML is slightly more accurate than IgPhyML.

**Fig. 4.**
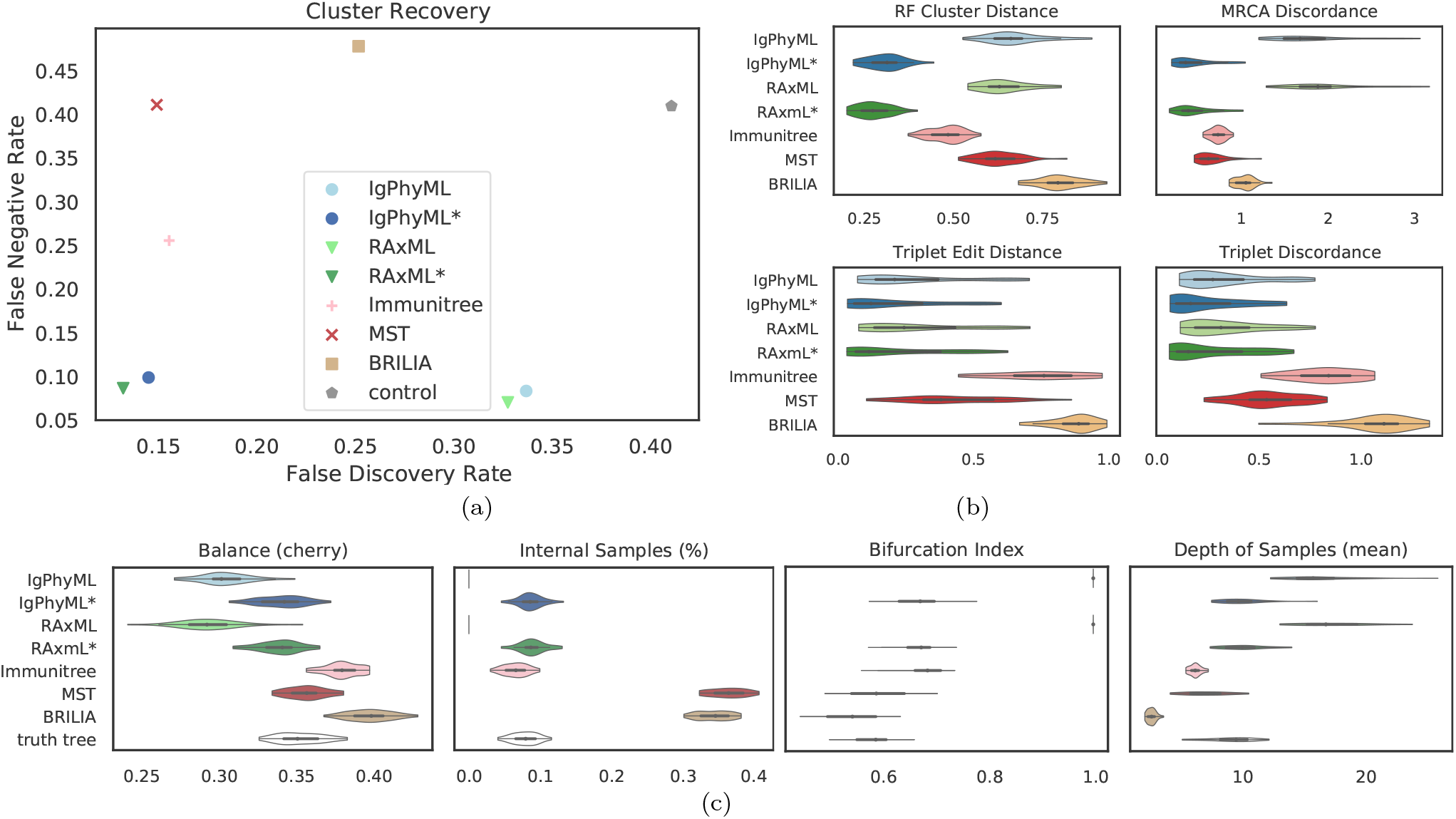
(a) False Discovery Rate (FDR) and False Negatie Rate (FNR) of various reconstruction methods on simulations under default conditions; (b) Normalized Robinson-Foulds cluster distance (RF), MRCA discordance (MD), triplet edit distance (TED), and triplet discordance (TD). (c) Properties of the estimated and true trees. Forresults excluding singeltons and the PD metric, see Fig. S2.

The MST-like methods have low FDR, coming close to zero-aware phylogeny-aware methods, but also have much higher FNR (25% or more). Immunitree (which uses Steiner trees) is substantially better than a simple MST in terms of FNR, but not in terms of FDR or triplet-based measures. These patterns largely follow the expectations: more resolved trees have lower FNRs whereas less resolved trees have lower FDRs. However, zero-aware phylogeny methods are able to obtain the best FDR and FNR and dominate other methods. BRILIA consistently has high error in our analyses. These patterns remain largely similar (but are magnified) when singletons are removed from the consideration (Fig. S2). The main exception is that when singletons are excluded, Immunitree is no longer the second best method according to the RF distance.

We next compare properties of the inferred trees and true trees (Figure 4c). BRILIA and MST put far too many labels at internal nodes (≈35% instead of ≈8%), while Immunitree and zero-aware phylogenetic trees are very close to the true tree in terms of percent internal samples. BRILIA and Immunitree over-estimate the tree balance, while phylogenetic trees under-estimate balance, especially before contracting low support branches. Conversely, phylogenetic methods over-estimate depth of samples while BRILIA, MST, and Immunitree underestimate the depth; zero-aware phylogenetic methods, however, produce trees that are very close to the true tree in sample depth. Phylogenetic methods, by definition, overestimate bifurcation index as 1; this overestimation is dramatically reduced but not fully eliminated by zero-aware phylogenetic methods and Immunitree. MST is quite close to the correct levels of bifurcation.

#### Varying selective pressure

The reconstructions methods are all impacted as selective pressure (*A*) changes, but some methods are more sensitive than others, and they are affected differently (Figs. 5ab). Zero-aware phylogenetic methods have the best accuracy across values of *A*. The ranking among other methods depends on the selective pressure such that phylogenetic methods become the worst when *A* is high and become the best when *A* is low. As *A* increases, error tends to increase for phylogenetic methods under all evaluation metrics except for the FNR; for example, the FDR of RAxML increases from 27% at the 1*/*4x level to 42% at the 2x level. In contrast, the error of Immunitree, MST, and BRILIA reduces with increased *A* according to FNR and RF. Zero-aware phylogenetic methods are relatively robust to the *A* and their error rates change only slightly across conditions. When singletons are removed from the metrics of comparison, patterns remain similar, though the impact of selective pressure becomes less pronounced (Fig. S3a).

**Fig. 5.**
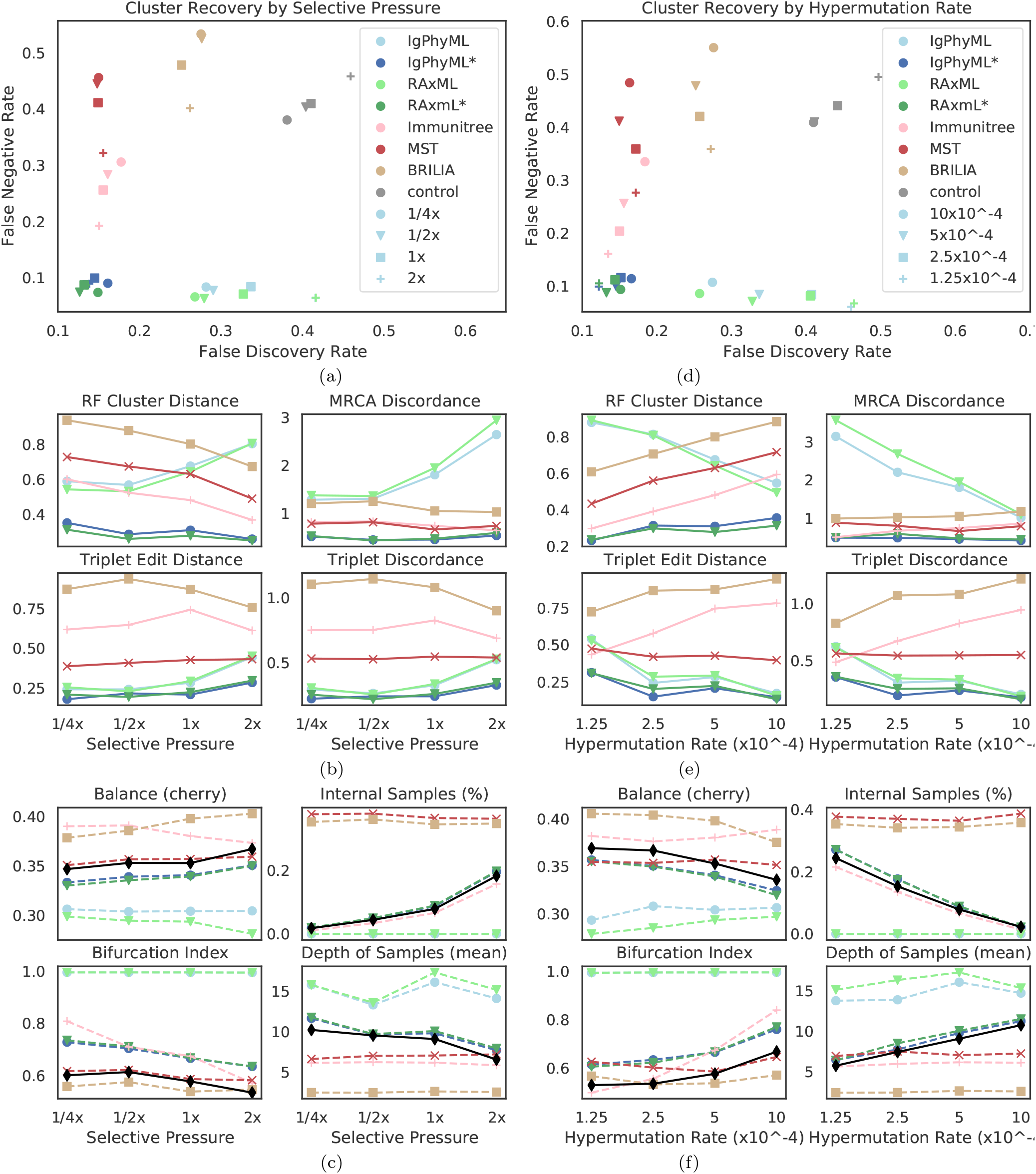
Impact of selective pressure *A* (a-c) and mutation rate *µ* (d-f) on tree inference error (a,b,d,e) and tree properties (c,f). We measure tree error by FDR and FNR (a,d), Robinson-Foulds cluster distance (RF), MRCA discorance (MD), triplet edit distance (TED), and triplet discordance (TD) (b,e). We show properties of true (black) and reconstructed trees (c,g). *µ* = 5 *×* 10^−5^ in (a-c) and *A* = 0.1 in (d-f), which are all default values.

The reason behind these patterns becomes more apparent once we consider changes in tree properties (Figs. 5c). As *A* increases, the fraction of internal samples tends to increase. This pattern can be explained: when selective pressure is high, cells with low affinity die off quickly, which results in shorter branch lengths. Since phylogenetic methods cannot put sequences on internal nodes, they have reduced accuracy. In contrast, IgPhyML*, RAxML*, and Immunitree are able to successfully assign sequences to internal branches; as a result, their percentage of internal samples match those of the true trees (Figs. 5c). Similarly, with increased *A*, the bifurcation index of the simulated tree tends to decrease, a pattern that is observed also in reconstructed trees from IgPhyML*, RAxML*, Immunitree, MST, and BRILIA. Again, phylogenetic trees, which produce binary trees, are unable to capture these patterns. As *A* increases, depth of sampled nodes of the simulated tree tends to decrease, a pattern matched by IgPhyML* and RAxML* but not other methods. Finally, when *A* is high, trees are shorter (i.e., accumulate less mutations) and more branches are single mutation (Fig. S4), both of which make phylogenetic inference more difficult. The reduced levels of depth, total change, and bifurcation make sense: higher pressure should result in fewer mutations needed to reach *M* because cells with unfavorable mutations are less likely to survive; this would produce shorter trees.

#### Varying rate of hypermutation

As the hypermutation rate (*µ*) increases, error decreases for normal Phylogenetic methods (IgPhyML and RAxML) according to most metrics but stays relatively stable for zero-aware methods (Fig. 5de). Increasing *µ* results in simulated trees that are marginally less balanced, are more bifurcating, have fewer internal node samples, and have a higher depth for sampled nodes (Fig. 5f). Thus, increasing *µ* generates trees more similar to what traditional phylogenetic methods assume. Zero-aware phylogenetic methods and Immunitree designate the right percentage of nodes as internal, but both are slightly more bifurcating than true trees (Fig. 5f). Overall, zero-aware phylogenetic methods are the most accurate across all values of *µ*.

#### Interplay between selective pressure and mutation rate

When we vary both *A* and *µ*, we observe that increasing mutation rate has similar effects on the error and tree properties as decreasing the selective pressure (Fig. 6). Reassuringly, error patterns observed when fixing one variable and changing the other are consistent with patterns when both variables are changed (Figs. 6 and S5). The most difficult condition for phylogenetic methods is low mutation rate and high selective pressure, where close to 70% of the branches include only a single mutation and bifurcation index is only 43%. However, zero-aware methods are impacted less in these conditions, and are in fact improved according to the RF metric (Fig. S5). In addition, we observe that antibody clonal trees become more phylogenetic-like – that is, more bifurcating (max: 0.74) and fewer internal samples (min: 20%) – with *µ* = 10^−3^ and *A* = 1*/*4x. Increasing the mutation rate or decreasing the selective pressure beyond these values leads to combinations where the infection could not be overcome.

**Fig. 6.**
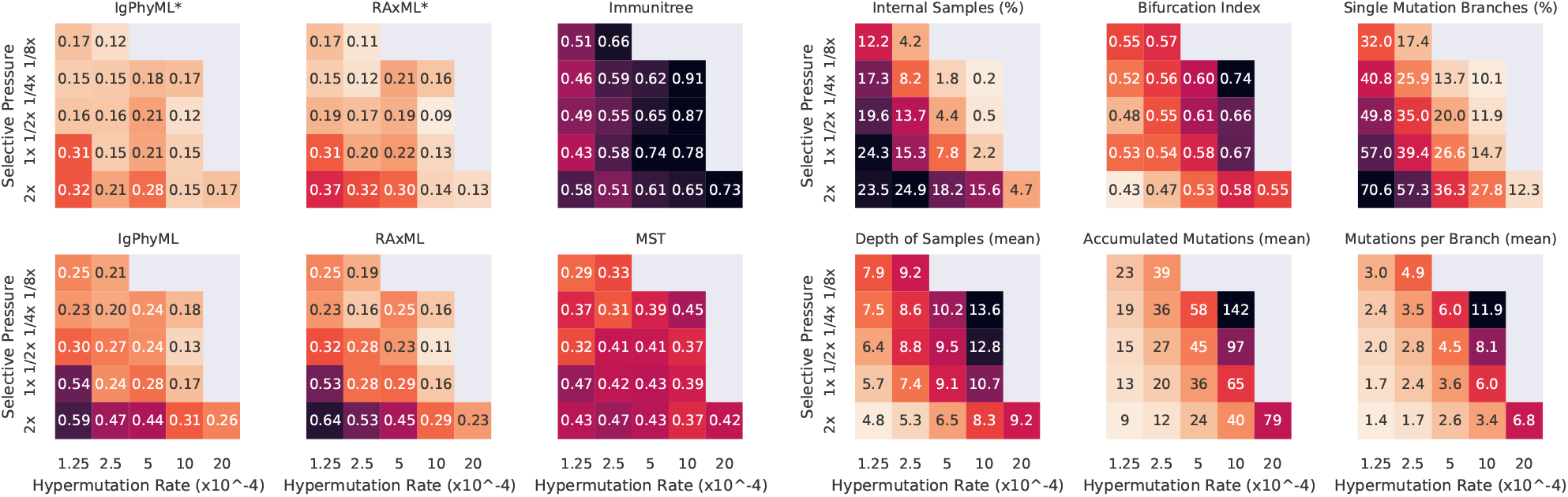
For varying levels of selective pressure (*A*), rate of hypermutation (*µ*), and all reconstruction methods except BRILIA, we show tree error measured by the triplet edit distance TED (left) and properties of the true tree (right). When the mutation rate is too high and selection pressure is to low, the simulation never ends, meaning that the total affinity needed to overcome the antigen is never reached; these conditions are missing from the figure. For other evaluation criteria see S5.

#### Other parameters

Beyond the main two parameters, we also studied changing six secondary parameters, most of which had relatively little impact on the results (Fig 7). As the weight of FRs regions in computing affinity (*w*_*f*_) increases, error tends to *slightly* increase for all methods under many evaluation metrics (Fig. S6). This pattern can be related to the slight increase in the number of single branch mutations and the reduction in the total number of substitutions across the tree. As germinal center capacity (*C*) increases, error increases or decreases slightly, depending on what measure is examined (Fig. S7). Increasing *C* tends to reduce internal samples of the simulated tree and single mutation branch and tends to increase mutations per branch. As memory cell life-time 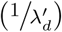 increases, error tends to increase for phylogenetic methods (Fig. S8), including IgPhyML* and RAxML*, which nevertheless continue to be the best methods. Plasma cells conversion rate (*ρ*_*p*_) (Fig. S9), rate of change in antibody target compared to antigen change (*κ*) (Fig. S10), and the threshold of total affinity for neutralization and stage change (*M*) (Fig. S11) have small and inconsistent impacts on tree inference error. In all conditions examined, IgPhyML* and RAxML* have the best accuracy (Fig 7).

**Fig. 7.**
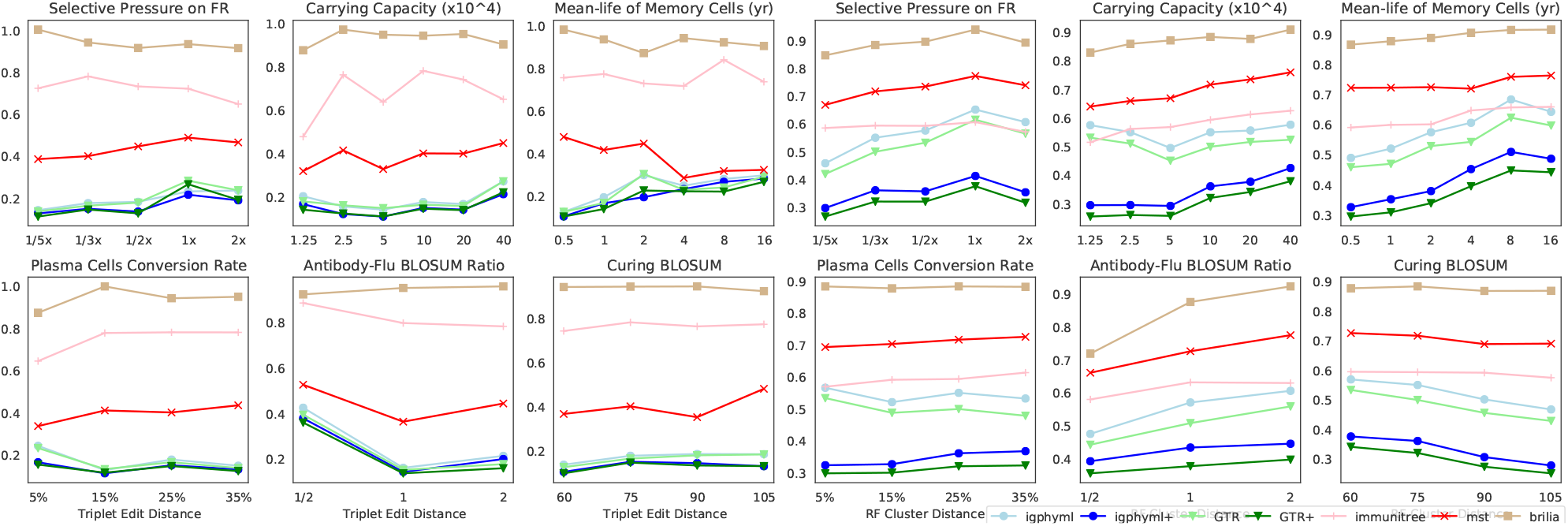
a) Triplet edit distances and b) RF cluster distances by selective pressure on framework region, carrying capacity, mean-life of memory cells, plasma cell conversion rate, antibody-flu blosum ratio (MARatio), stage change threshold (M).

## Discussion

### Implications for reconstructing antibody evolution

Our study partially confirms that phylogenetic methods need to change for inferring antibody clonal trees with high accuracy. Depending on the simulation condition, 1% to 20% of sampled sequences belonged to internal nodes, and the true trees are only 60% to 70% bifurcating. We observed that results of phylogenetic inference using ML, taken at face value, can have low accuracy. However, we also showed that ML phylogenetic methods, with a very simple adjustment, can outperform the alternative methods based on Steiner trees and spanning trees. The simple adjustment we applied was to contract branches with length lower than a fixed constant. We selected this constant using a rule-of-thumb based on the length of the sequences; however, statistical tests of whether a zero branch length null hypothesis can be rejected exist (Jackman *et al*., 1999; Walsh *et al*., 1999; Goldman *et al*., 2000) and are fast (Anisimova *et al*., 2006) and could be used *in lieu* of our simple heuristic. Moreover, our work implies that phylogenetic methods that try to naturally model zero branch length (e.g., Lewis *et al*., 2005) are also promising. In particular, the adaptive LASSO method of Zhang *et al*. (2020) seems suitable for inferring antibody evolution and should be put to test once available as part of a software package.

Despite the higher accuracy of zero-aware phylogenetic methods compared to the available alternatives, we note that there is still substantial error. Under the default condition, 90% of clusters of the true tree were recovered but about 15% of the recovered clusters were incorrect. In particular, the discrepancy between FNR and FDR is due to the fact that the inferred trees are somewhat more bifurcating than true trees (e.g., ≈70% versus 60% in the default condition). Thus, while contracting some super-short branches has been helpful in increasing accuracy, our zero-aware phylogenetic trees are still biased towards too much resolution. It is possible that better Steiner-based methods that incorporate more advanced models of sequence evolution can solve this shortcoming.

### Implications for evaluation criteria

The ranking of reconstruction methods can change based on which of the ten evaluation criteria we choose, and these rankings only partially correlate (Fig. S12). Most interestingly, FDR and FNR are weakly *anti*-correlated (mean Spearman’s rank correlation coefficient across all tests *ρ* = −0.12), though excluding singletons changes this patterns. Thus, false positive and false negative errors can a paint contradictory picture, especially when singletons are included. RF distance, which combines both aspects, correlates moderately with both FDR (*ρ* = 0.5) and FNR (*ρ* = 0.57). The triplet-based metrics strongly agree with each other (*ρ* = 0.97) and are mostly compatible with the RF distance (*ρ* ≈ 0.75), but are less similar to MD and PD metrics (*ρ* ⩽ 0.52). Consistent with the observation that triplet metrics penalize false negatives more than false positives, they agree more strongly with FNR than FDR (*ρ* = 0.65 vs 0.26). MD and PD are very similar to each other (*ρ* = 0.96), have no correlation to FNR (*ρ* 0.05), but have moderately high correlation to FDR (*ρ* = 0.71). Finally, we notice that singletons can matter: while FNR and FNR* are highly correlated (*ρ* = 0.94), RF correlates with RF* less strongly (*ρ* = 0.71), and FDR correlates with FDR* only moderately (*ρ* = 0.61).

The choice of the metric should depend on downstream application of the clonal tree. While zero-aware phylogenetic methods are judged to be dramatically better than normal phylogenetic methods based on most criteria, they are only slightly better according to the triplet-based criteria. The triplet metrics do not penalize trees heavily if they are more resolved than the true tree or if they move internal nodes to leaves. Thus, when downstream usage is robust to extra resolution and extra terminal edges, triplet metrics offer a good way to measure topological accuracy. On the other extreme, PD and MD are very sensitive to the tree resolution and internal placement, so much so that they often evaluate inferred phylogenetic trees to be much worse than random trees (Fig. S5) because these trees generate fully resolved trees and put samples at leaves. Thus, we don’t find PD and MD to be reliable metrics of *topological* accuracy. RF distance is in between: it penalizes extra resolution more than triplet metrics but less than path-based metrics. It does distinguish zero-aware and phylogenetic methods but rarely evaluates any methods to be worse than random (Fig. S5). Overall, dividing the observed error along two (potentially contradictory) axes such as FNR and FDR is recommended because this evaluation provides more insight into reasons behind error.

### Comparison to outer simulation models

Several simulation tools capable of benchmarking reconstruction methods have been recently developed. Some of these tools are not comparable to our effort because of various limitations. The recent immuneSIM by Weber *et al*. (2020) generates mutations but does not model the clonal tree or the selection process. Methods of Amitai *et al*. (2017) and Reshetova *et al*. (2017) are based on the two-step simulation paradigm and only generate clonal trees under selection, leaving sequences generation to other methods. The most relevant method to ours are bcr-phylo by Davidsen and Matsen (2018) and gcdynamics by Childs *et al*. (2015), which simulate clonal trees of antibody-coding sequences under AM. Both bcr-phylo and gcdynamics have similarities and differences to our method (Table 6). For example, they both support multiple targets but only one round of simulations. Although our model is capable of multiple targets, for simplicity, DIMSIM uses one target per round of infection. However, the advantage of DIMSIM is that, unlike the two other methods that only simulate activated cells, it also simulates memory cells; as a result, it can simulate multiple rounds of infection by an evolving antigen with changing targets while considering memory built from previous infections. Moreover, DIMSIM simulates in continuous time whereas the other tools simulate under discrete generations. All three methods use sequences to define affinity, albeit differently: DIMSIM using BLOSUM distance, brc-phylo using hamming distance, and gcdynamics using random energy landscape. A main feature of DISMSIM is that its birth/death rates are polynomial fractions of individual and total affinity; this choice enables it to speed up the simulation, allowing it to scale up to millions of cells, unlike the other two methods. Advantages of the other tools include the fact that only brc-phylo simulates isotype switching and only gcdynamics distinguishes intra- versus inter- germinal center competitions.

**Table 6.** A comparison of Most relevant tools for AM simulation.

|  | DIMSIM<br>this paper | bcr-phylo<br>Davidsen and Matsen (2018) | gcdynamics<br>Childs *et al.* (2015) |
| --- | --- | --- | --- |
| Targets | Single-target (per round) | Multi-target (1 round) | Multi-target (1 round) |
| Rounds | Yes | No | No |
| Affinity | BLOSUM distance | Hamming distance | Random energy landscape |
| Mutation | Updated Yaari *et al.* (2013) | Yaari *et al.* (2013) | i.i.d |
| Scalability | Up to millions of cells | Thousands of cells | Thousands of cells |
| Cell type | Activated & Memory | Activated | Activated |
| Germinal Centers | Combined (single) | Combined (single) | Multiple (in competition) |
| Time | Continuous | Discrete generations | Discrete generations |
| Isotype | No | Yes | No |
| Birth/Death rate | Polynomial fraction of<br>individual and total affinity | Neutral: independent of total affinity<br>Kinetic: function of affinities | A function of affinity |

### Limitations of the study

Our study has limitations that should be kept in mind.

In our simulations, we did not add errors to sequence data used as input to clonal tree reconstruction methods. Real Rep-Seq samples undergo extensive PCR and thus might contain both sequencing and amplification errors. We assumed that error elimination is already performed (to perfection) prior to reconstruction using existing methods (e.g., Vander Heiden *et al*., 2014; Safonova *et al*., 2015; Bolotin *et al*., 2015; Shlemov *et al*., 2017). We also simulated only substitution SHMs but no insertions and deletions. We note that, in these shortcoming, our study is not different from most phylogenetics simulations that also fail to incorporate indels and many forms of errors in input, such as alignment error, orthology error, and assembly error. Nevertheless, the impact of the error on various methods and the overall accuracy should be tested in future work. Similarly, the efficacy of methods that simultaneously filter errors and build clonal trees (e.g., Safonova and Pevzner, 2019; Lee *et al*., 2017) should be subject of future research.

In our AM model, we had to adopt several arbitrary assumptions in order to simulate the selective pressure. For example, absent of a good model of receptor binding, we assumed the affinity grows gradually as the AA sequence becomes more similar to the target sequence (i.e., the best possible antibody for an antigen). The idea that AM occurs by mutational diffusion along one or more preferred paths in the genotype space has been supported by Kepler *et al*. (2014). Nevertheless, our i.i.d model is certainly a simplification without a clear empirical support. Moreover, we assumed the existence a target antibody sequence. The literature has increasingly documented highly convergent immune responses to the same epitope across individuals and conditions (Henry Dunand and Wilson, 2015; Robbiani *et al*., 2020). This observation gives us reason to think the existence of target sequences is not a bad assumption; nevertheless, the choice of a *single* target may not be realistic. To model the change in the target as the viruses evolve across seasons, we chose targets with evolutionary divergence levels that mimic divergence levels of the antigen, albeit with some scaling factor. While we believe this choice is sensible, again, we have no evidence to back up this model on empirical grounds. It is conceivable that two antigens with high evolutionary distance are neutralized by similar antibodies, or that, antigens that are very similar require very distant antibodies. Finally, our 5-mer mutation model, while based on the empirical model of Yaari *et al*. (2013), still fails to capture some of the complexities of the real antibody evolution. For example, we concentrated substitutions on the CDR region, but other regions are known to also accumulate mutations (Safonova and Pevzner, 2019; Kirik *et al*., 2017; Ovchinnikov *et al*., 2018). Other B cell specific models (e.g., Elhanati *et al*., 2015) including those that seek to tease out the effects of selection from background mutations (e.g., McCoy *et al*., 2015) and per-position mutability models (Kepler *et al*., 2014) can be incorporated in the future.

For all these shortcomings in modelling, we offer several responses. The framework is designed to be flexible and can easily incorporate more complex models if a better understanding of processes behind antibody-antigen affinity is achieved (e.g., Luo and Perelson, 2015) and is formalized in mathematical models. Thus, our work should be considered a first step that will enable better modeling in future. We also remind the reader that our objective was to simulate so that we can benchmark various tools for reconstructing clonal trees. Thus, as long as our modelling choices did not distort the comparison of methods, some model misspecification can be tolerated. We observed that the choice of the best method was not sensitive to many parameter choices.

Beyond model simplifications, we also chose to simulate parts of the complex immune system response, but not others. For example, we simulated one clonal lineage involved in an immune response. As such, we ignored the important VDJ recombination step and sought to simply simulate a VDJ recombinant that is effective in fighting a specific antigen. Even then, we simulated only one clonal lineage at a time, a limitation that can be easily lifted in the future by starting from multiple root sequences with different VDJ settings and assigning to each a different target sequence. Note that our tool can be easily combined with methods of simulating VDJ recombination such as IGoR (Marcou *et al*., 2018). Neither did we simulate light chains, which are often not captured in Rep-Seq sequencing data, but we note that extending the methodology to light chains, given better understanding of their evolution, will be possible.

Finally, while we tested several reconstruction methods, we were not able to test others. We are unable to install SAMM (Davidsen and Matsen, 2018) and GCtree (DeWitt *et al*., 2018) due to their dependencies, and we were unable to find an implementation of the IgTree (Barak *et al*., 2008) method.

### Applications of the framework

The framework we designed for simulation of clonal trees can be extended for simulating other forms of micro-evolutionary scenarios. While the current implementation is geared towards AM simulations, our proposed algorithm enables forward-time simulation of very large numbers of entities under models that allow dependence between sequences and rates of birth, death, or transformation. The ability to simulate a very large number of entities combined with rates that change with properties of entities give use the necessary ingredients to simulate under complex models of evolution that consider selective pressure. Thus, our framework can be adopted for other forms of micro-evolutionary simulation such as the evolution of a virus within a host and accumulation of SHMs in tumor evolution. Such a possibility would become most intriguing if it can also model co-evolution of different types of entities (e.g., antibodies and viruses). While we did not simulate co-evolution here, we believe the framework is capable of performing such simulations by simply creating entity types (just like we had cell types) and making the BDT rates a function of properties across different cell types. Another promising direction for extensions of this work is to integrate the sequence evolutionary models with network-based disease transmissions models (e.g., Ratmann *et al*., 2017; Moshiri *et al*., 2019) to enable more accurate simulations of disease spread and evolution.

## Availabilty

DIMSIM simulation framework and relate code is publicly available at https://github.com/chaoszhang/immunosimulator. All the data are available at https://github.com/chaoszhang/DIMSIM-data.

## Acknoeledgements

We would like to thank Pavel A. Pevzner, Li-Fan Lu, and Jiawang Nie for insightful discussions on clonal tree reconstruction, affinity maturation, and optimization methods. This work was supported by National Science Foundation (NSF) grant III-1845967. Computations were performed on the San Diego Supercomputer Center (SDSC) through XSEDE allocations, which is supported by the NSF grant ACI-1053575.

## APPENDIX

### Supplementary Materials

#### Supplementary Methods

##### Derivation of Equation (1)

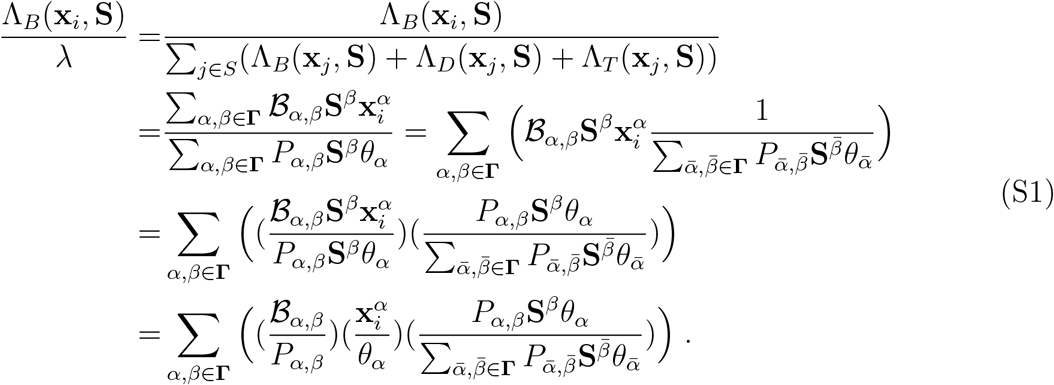

##### Somatic hypermutagenesis frequency models for **K**^5^ and f

Our model is based on an empirical frequency **K**^5^(*s, s*_1_, *s*_2_, *s*_3_, *s*_4_, *s*_5_) matrix that counts the number of times 5-mer (*s*_1_, *s*_2_, *s*_3_, *s*_4_, *s*_5_) converts to (*s*_1_, *s*_2_, *s, s*_4_, *s*_5_) in one cycle of cell division during hyper-mutation. Given the matrix, we define

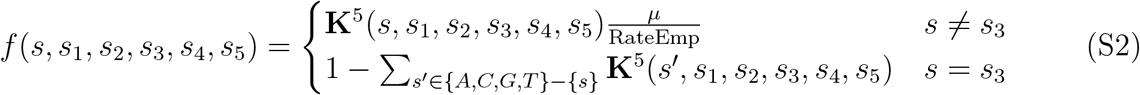

where

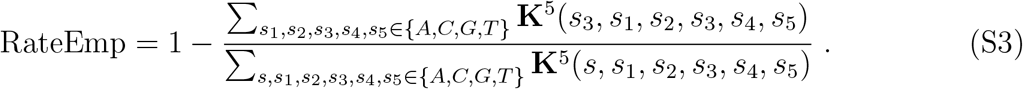

Somatic hypermutagenesis of antibodies is the result of activation-induced deaminase (AID) enzyme activity that changes a random C:G base into a U:G base in B cell DNA. U:G mismatch can be repaired using UDG (uracil-DNA glycosylase) or MMR (DNA mismatch repair) machinery that forms diversity of hypermutations (Peled *et al*., 2008). Certain biological mechanisms of SHM occurrences were studied extensively. For example, Rogozin and Kolchanov (1992) observed specific hot/cold-spot DNA motifs for SHMs in immunoglobulin genes. Particularly, WRCY/RGYW where W = *{*A, T*}*, Y = *{*C, T*}*, R = *{*G, A*}* and later predicted more general WRCH/DGYW with H = *{*A, C, T*}* and D = *{*A, G, T*}* motifs are hot-spots for SHMs caused by weak hydrogen-bounds (Rogozin and Diaz, 2004). SYC/GRS (S = C, G) is a cold-spot motif caused by strong hydrogen-bounds (Bransteitter *et al*., 2004). The locality of AID enzyme activity has been emphasized. (Smith *et al*., 1996; Shapiro *et al*., 2003).

To simulate SHM we modified a model proposed by Yaari *et al*. (2013). The model extends the notion of hot/cold-spots and suggests that a certain hierarchy of mutabilities exists following Smith *et al*. (1996) and Shapiro *et al*. (2003). The model is based on the mutability of a central base in each 5-mer of an antibody heavy chain and consists of two parts: a targeting model identifying if a mutation occurs in the variable part of an antibody and a substitution model providing an insight into what is this mutation. In order to avoid selection bias, the authors considered 5-mers where only synonymous substitutions of the central base are possible and inferred probabilities for other 5-mers. Unfortunately, synonymous substitutions constitute only a fraction of possible mutations. To overcome this issue Yaari *et al*. (2013) proposed a special inference method to estimate parameters for the rest of 5-mers. Parameters for targeting and substitution models were inferred for 468 and 740 5-mers, respectively. However, the accuracy of this procedure was shown to be sub-optimal (Yaari *et al*., 2013, Table 2). Additionally, some of the datasets that were used to estimate the parameters are derived from an error-prone 454 sequencing technology.

We re-estimated the parameters of this model and considered all 5-mers without limiting our scope to synonymous mutations. We also utilized three up-to-date repertoire sequencing datasets (all data was produced using the Illumina MiSeq platform): *i*) PRJNA349143. Time series of three individuals during influenza vaccination, both before and after vaccination. *ii*) PRJNA395083. Bulk unsorted PBMC from peripheral blood of several healthy donors. *iii*) A dataset of paired end sequences, added to increase power. While the last dataset we used is not publicly available, we make the resulting k-mer model available publicly at https://github.com/chaoszhang/immunosimulator/blob/master/kmerFreq.txt.

From each dataset we obtained a matrix of the size 1024 *×* 4, where each row corresponds to a distinct 5-mer and contains *# nonmutated occurrences* of this 5-mer and three possible *# nucleotide substitution occurrences*. To calculate this matrix for a given dataset, we found the closest V gene for every read and record the number of observed 5-mers in the gene and their corresponding mutated copies across the read. For any 5-mer *K*, the corresponding row of a constructed matrix can be viewed simultaneously as a value of *Binomial* and *Multinomial* distributions. *Binomial* distribution represents the number of occurred mutations among all occurrences of the 5-mer *K*, while *Multinomial* distribution indicates the number of mutations to specific bases among all occurred mutations. The parameters of these distributions indicate the mutability and substitution profiles for each 5-mer *K*. The 5-mer frequencies were combined across all these datasets to obtain the final matrix, available at https://github.com/chaoszhang/immunosimulator/blob/master/kmerFreq.txt.

##### Default parameters

Here we provide the actual default values used for several parameters that did not fit in Table 1.

*BLOSUM*. The BLOSUM matrix table (Table S1) is obtained from ftp://ftp.ncbi.nih.gov/blast/matrices/BLOSUM100.

**Table S1.**
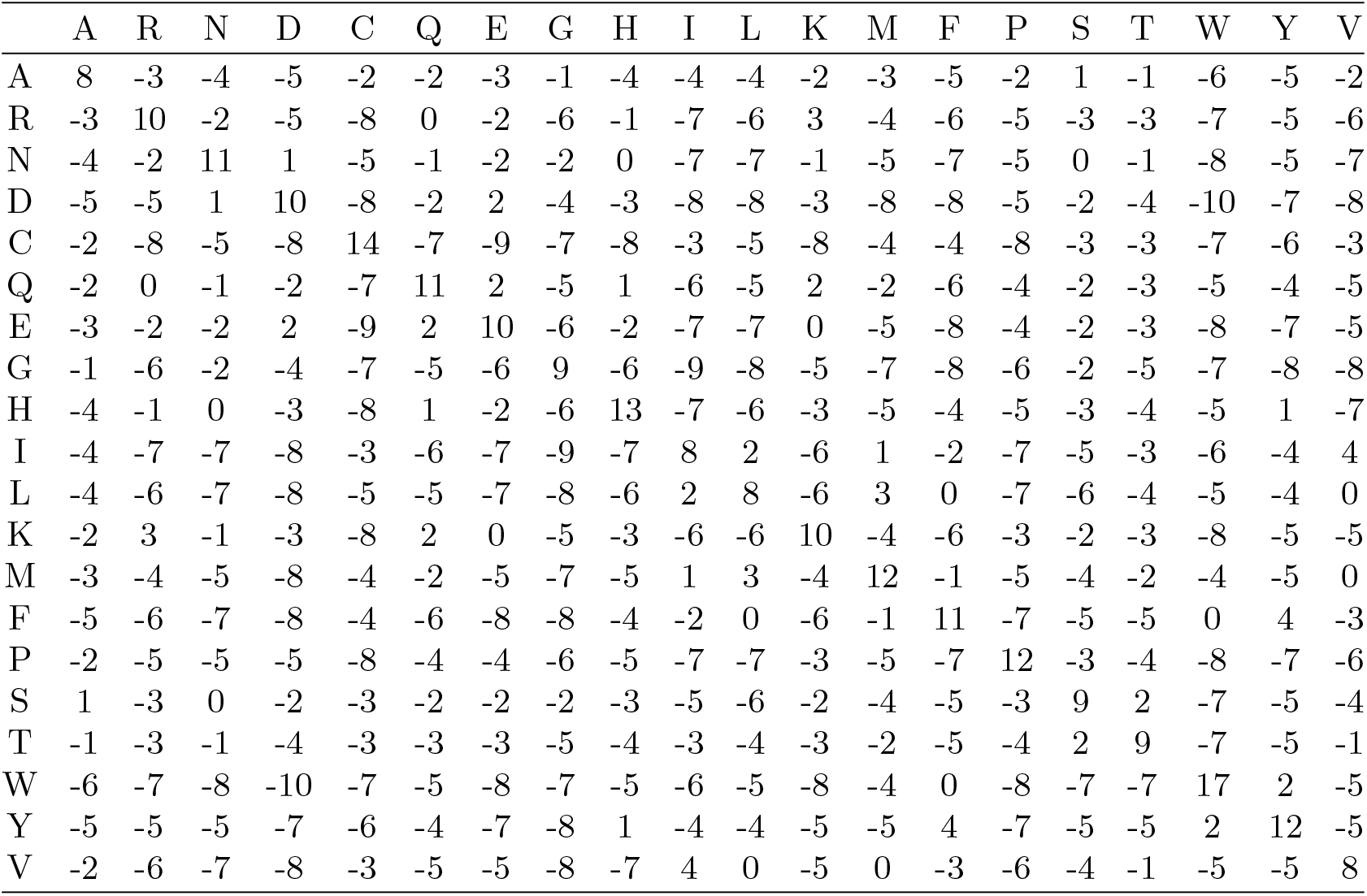
BLOSUM table

|  | A | R | N | D | C | Q | E | G | H | I | L | K | M | F | P | S | T | W | Y | V |
| --- | --- | --- | --- | --- | --- | --- | --- | --- | --- | --- | --- | --- | --- | --- | --- | --- | --- | --- | --- | --- |
| A | 8 | -3 | -4 | -5 | -2 | -2 | -3 | -1 | -4 | -4 | -4 | -2 | -3 | -5 | -2 | 1 | -1 | -6 | -5 | -2 |
| R | -3 | 10 | -2 | -5 | -8 | 0 | -2 | -6 | -1 | -7 | -6 | 3 | -4 | -6 | -5 | -3 | -3 | -7 | -5 | -6 |
| N | -4 | -2 | 11 | 1 | -5 | -1 | -2 | -2 | 0 | -7 | -7 | -1 | -5 | -7 | -5 | 0 | -1 | -8 | -5 | -7 |
| D | -5 | -5 | 1 | 10 | -8 | -2 | 2 | -4 | -3 | -8 | -8 | -3 | -8 | -8 | -5 | -2 | -4 | -10 | -7 | -8 |
| C | -2 | -8 | -5 | -8 | 14 | -7 | -9 | -7 | -8 | -3 | -5 | -8 | -4 | -4 | -8 | -3 | -3 | -7 | -6 | -3 |
| Q | -2 | 0 | -1 | -2 | -7 | 11 | 2 | -5 | 1 | -6 | -5 | 2 | -2 | -6 | -4 | -2 | -3 | -5 | -4 | -5 |
| E | -3 | -2 | -2 | 2 | -9 | 2 | 10 | -6 | -2 | -7 | -7 | 0 | -5 | -8 | -4 | -2 | -3 | -8 | -7 | -5 |
| G | -1 | -6 | -2 | -4 | -7 | -5 | -6 | 9 | -6 | -9 | -8 | -5 | -7 | -8 | -6 | -2 | -5 | -7 | -8 | -8 |
| H | -4 | -1 | 0 | -3 | -8 | 1 | -2 | -6 | 13 | -7 | -6 | -3 | -5 | -4 | -5 | -3 | -4 | -5 | 1 | -7 |
| I | -4 | -7 | -7 | -8 | -3 | -6 | -7 | -9 | -7 | 8 | 2 | -6 | 1 | -2 | -7 | -5 | -3 | -6 | -4 | 4 |
| L | -4 | -6 | -7 | -8 | -5 | -5 | -7 | -8 | -6 | 2 | 8 | -6 | 3 | 0 | -7 | -6 | -4 | -5 | -4 | 0 |
| K | -2 | 3 | -1 | -3 | -8 | 2 | 0 | -5 | -3 | -6 | -6 | 10 | -4 | -6 | -3 | -2 | -3 | -8 | -5 | -5 |
| M | -3 | -4 | -5 | -8 | -4 | -2 | -5 | -7 | -5 | 1 | 3 | -4 | 12 | -1 | -5 | -4 | -2 | -4 | -5 | 0 |
| F | -5 | -6 | -7 | -8 | -4 | -6 | -8 | -8 | -4 | -2 | 0 | -6 | -1 | 11 | -7 | -5 | -5 | 0 | 4 | -3 |
| P | -2 | -5 | -5 | -5 | -8 | -4 | -4 | -6 | -5 | -7 | -7 | -3 | -5 | -7 | 12 | -3 | -4 | -8 | -7 | -6 |
| S | 1 | -3 | 0 | -2 | -3 | -2 | -2 | -2 | -3 | -5 | -6 | -2 | -4 | -5 | -3 | 9 | 2 | -7 | -5 | -4 |
| T | -1 | -3 | -1 | -4 | -3 | -3 | -3 | -5 | -4 | -3 | -4 | -3 | -2 | -5 | -4 | 2 | 9 | -7 | -5 | -1 |
| W | -6 | -7 | -8 | -10 | -7 | -5 | -8 | -7 | -5 | -6 | -5 | -8 | -4 | 0 | -8 | -7 | -7 | 17 | 2 | -5 |
| Y | -5 | -5 | -5 | -7 | -6 | -4 | -7 | -8 | 1 | -4 | -4 | -5 | -5 | 4 | -7 | -5 | -5 | 2 | 12 | -5 |
| V | -2 | -6 | -7 | -8 | -3 | -5 | -5 | -8 | -7 | 4 | 0 | -5 | 0 | -3 | -6 | -4 | -1 | -5 | -5 | 8 |

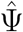 *and ζ*_0_. The starting sequence 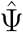 is set to be CAGGTGCAGCTGCAGGAGTCGGGCCCAGG ACTGGTGAAGCCTTCACAGACCCTGTCCCTCACCTGCACTGTCTCTGGTGGCTCCATCAGCAGTGGTGGTTACTA CTGGGCTGGATCCGCCAGCACCCAGGGAAGGGCCTGGAGTGGATTGGGTACATCTATTACAGTGGGAGCACCTA CTACAACCCGTCCCTCAAGAGTCGAGTTACCATATCAGTAGACACGTCTAAGAACCAGTTCTCCCTGAAGCTGAG CTCTGTGACTGCCGCGGACACGGCCGTGTATTACTGTGCGAGAGCGCGCGTCAATAGGGATATTGCGTACGGCAA CTGGTTCGACCCCTGGGGCCAGGGGACCCTGGTCACCGTCTCCTCA and thus *ζ*_0_ is QVQLQESGPGLVKPSQT LSLTCTVSGGSISSGGYYWSWIRQHPGKGLEWIGYIYYSGSTYYNPSLKSRVTISVDTSKNQFSLKLSSVTAADT AVYYCARARVNRDIAYGNWFDPWGQGTLVTVSS.

*η*_*i*_, *ζ*_*i*_, *and t*_*i*_. Are given in Table S2.

**Table S2.**
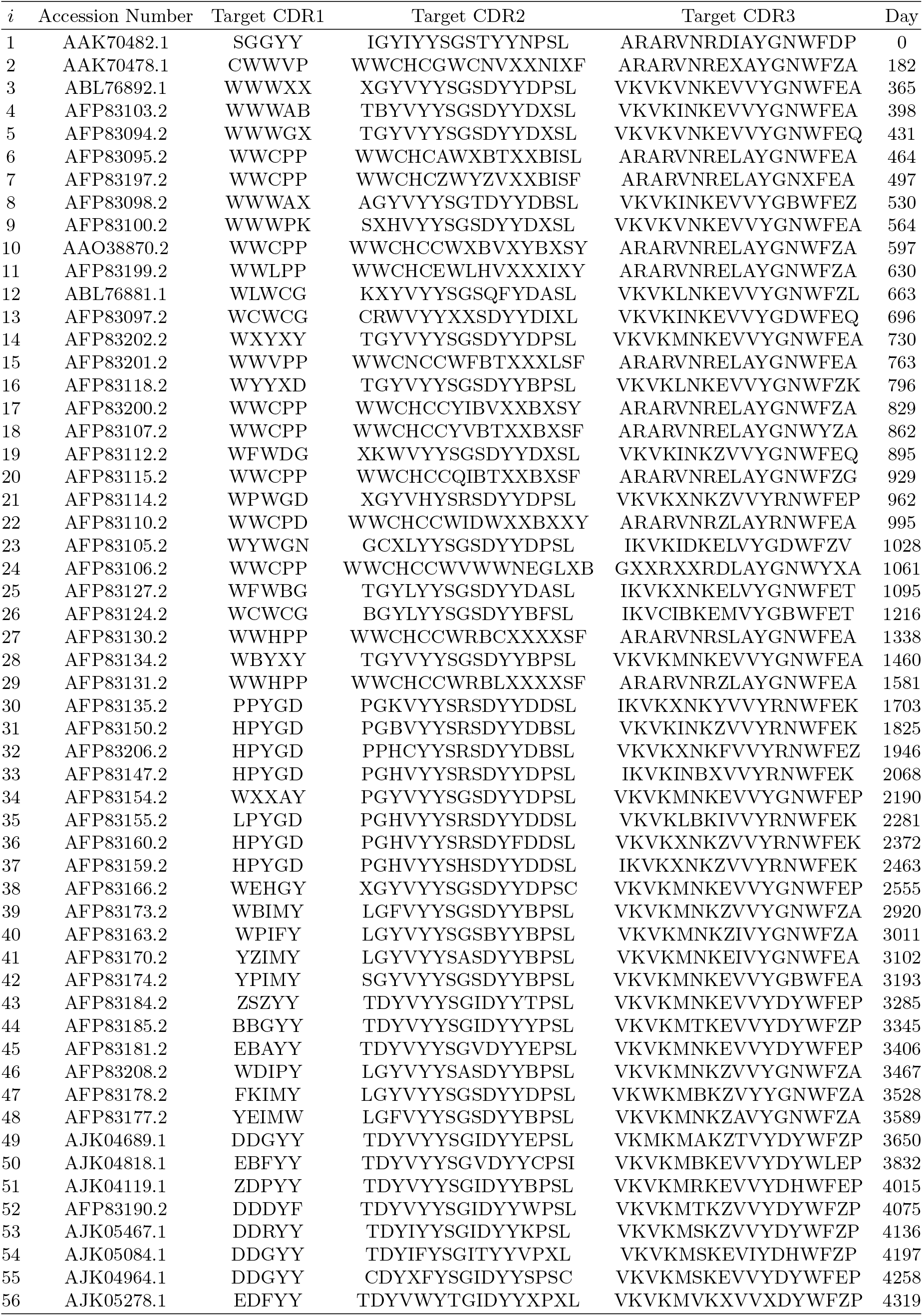
Flu accession number, CDRs of target sequences, and starting day of infection

#### Supplementary Tables and Figures

**Table S3.** Birth, death, and transformation rate functions as polynomials.

| Rate functions | Infected stage | Dormant stage |
| --- | --- | --- |
| $\Lambda_B(\mathbf{x}_i, \mathbf{S})$ | $\lambda_b g_i$ | 0 |
| $\Lambda_D(\mathbf{x}_i, \mathbf{S})$ | $\frac{\lambda_b(1-\rho_p-\rho_m)}{C}(\frac{g_i}{a_i})\sigma + (\rho_p\lambda_b - \lambda'_d)g_i + \lambda'_d$ | $(\lambda_d - \lambda'_d)g_i + \lambda'_d$ |
| $\Lambda_T(\mathbf{x}_i, \mathbf{S})$ | $t_i$ | 0 |

**Fig. S1.**
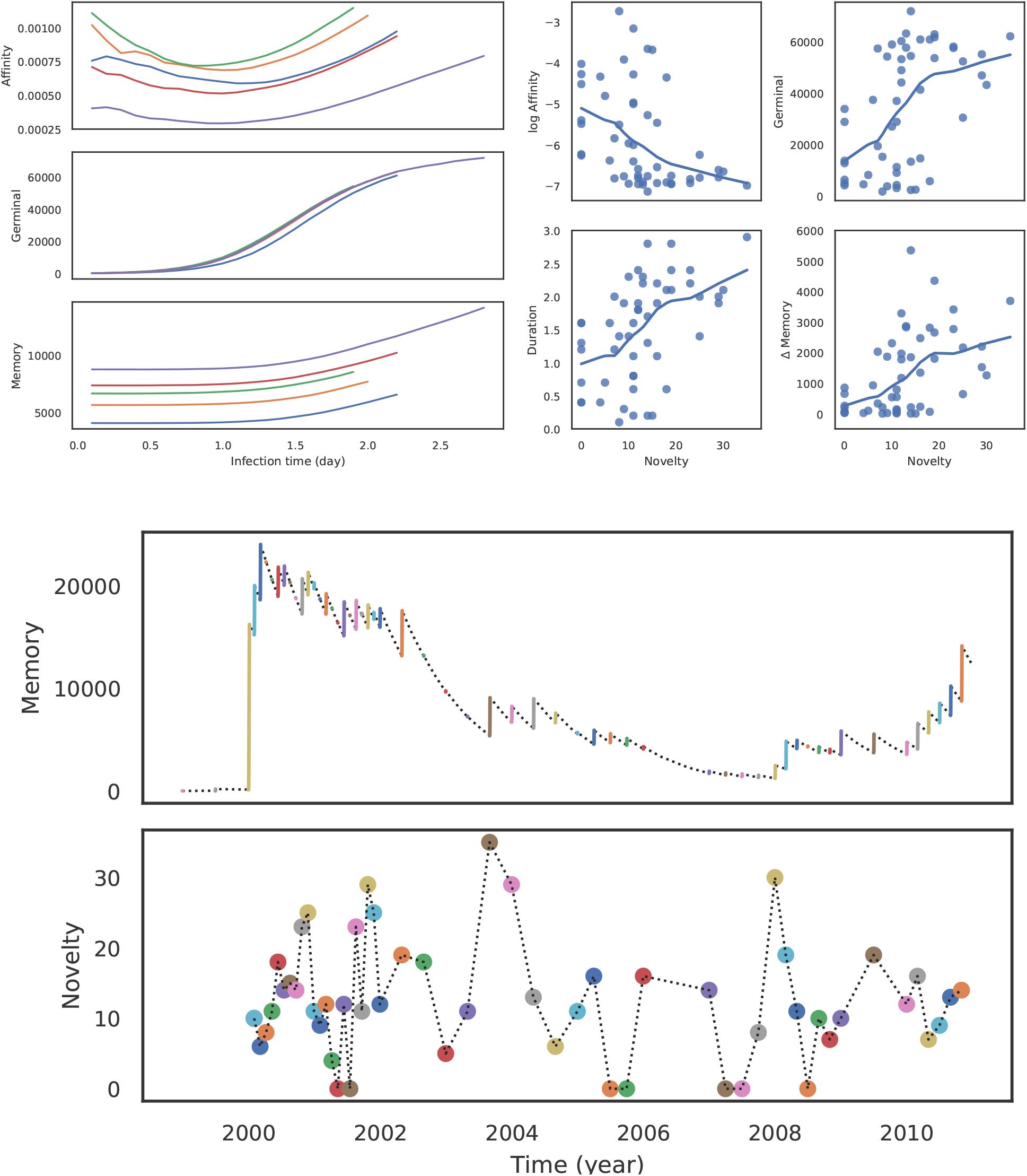
a) Log average affinity of activated cells to current infection target at the end of the infection, the number of activated cells at the end of the infection, and the duration of infection by novelty of the target of one simulation under default conditions, showing the last five rounds as examples. b) Average affinity of activated cells to current infection target, the number of activated cells, and the number of memory cells by time after infection starts for the last five infections of one simulation under default conditions. Lines are fitted using the LOWESS (locally weighted scatterplot smoothing) algorithm. c) Number of memory cells and novelty of infections by time. Dormant stages are indicated by dotted lines.

**Fig. S2.**
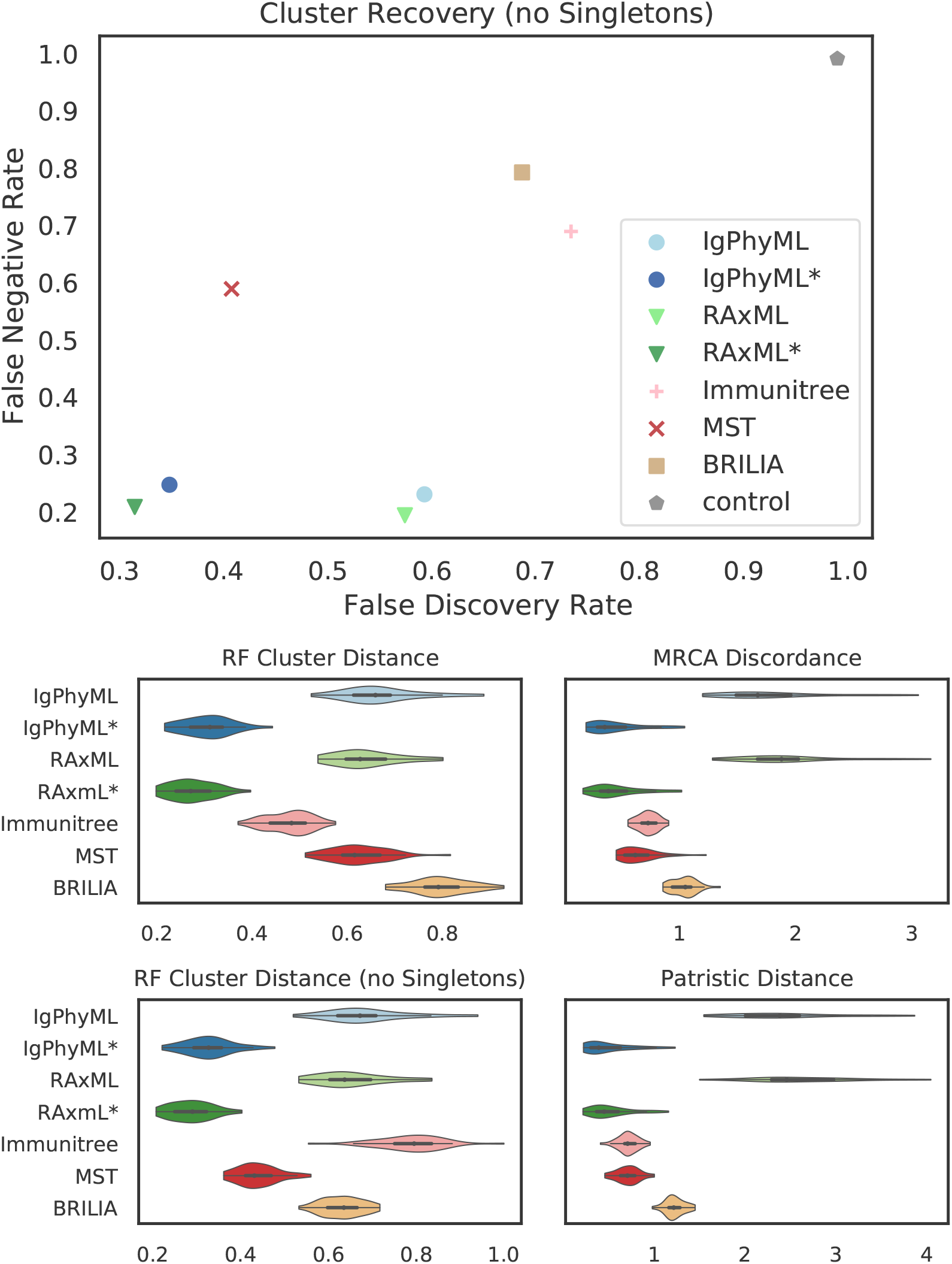
Top: FNR* and FPR* rates excluding singletons by reconstruction methods on simulations under default conditions; Bottom: Normalized Robinson-Foulds cluster distance with and without singletons (RF and RF *), MD and PD.

**Fig. S3.**
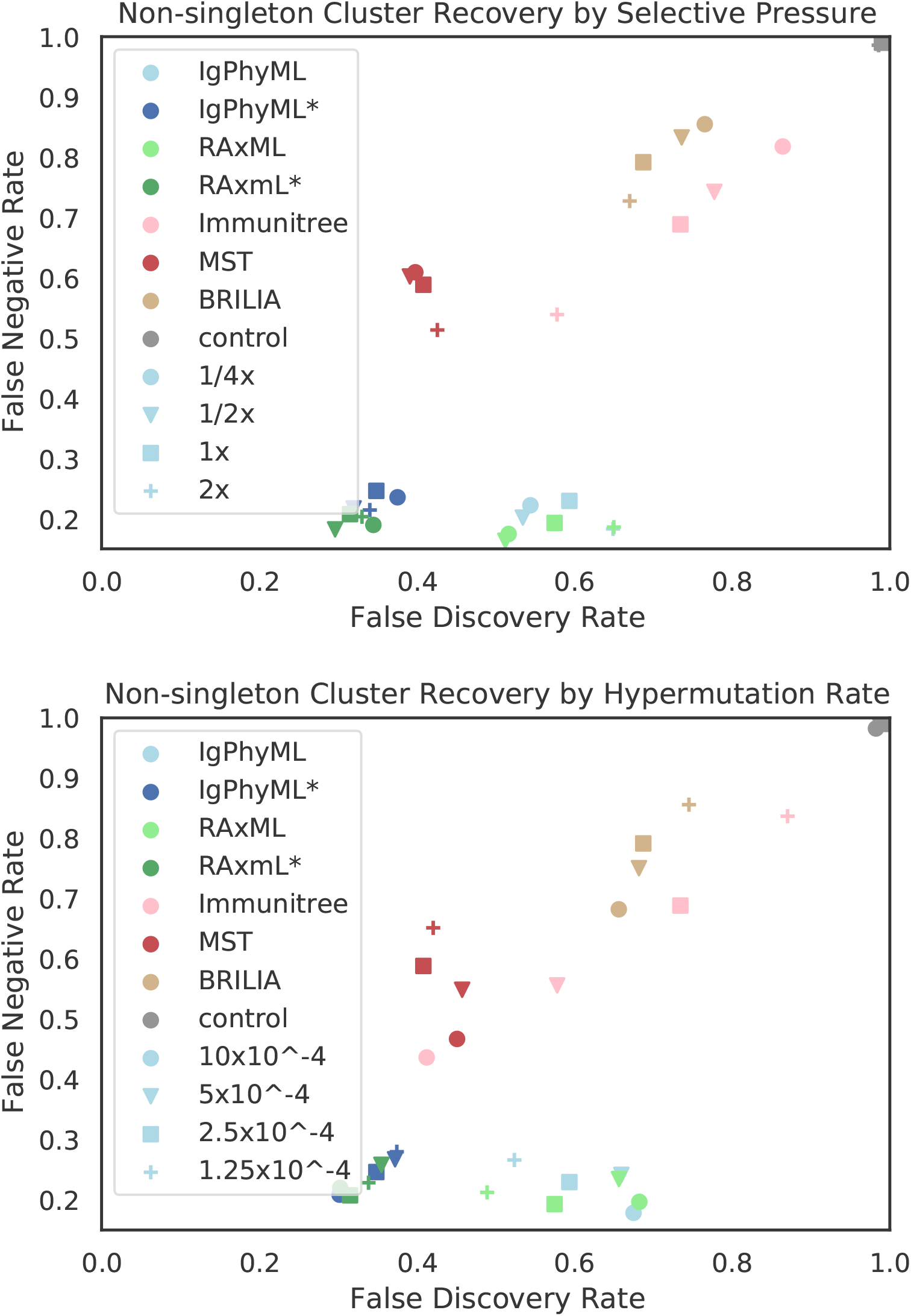
Impact of selective pressure *A* (a) and mutation rate *µ* (b) on tree inference error by FDR* and FNR*.

**Fig. S4.**
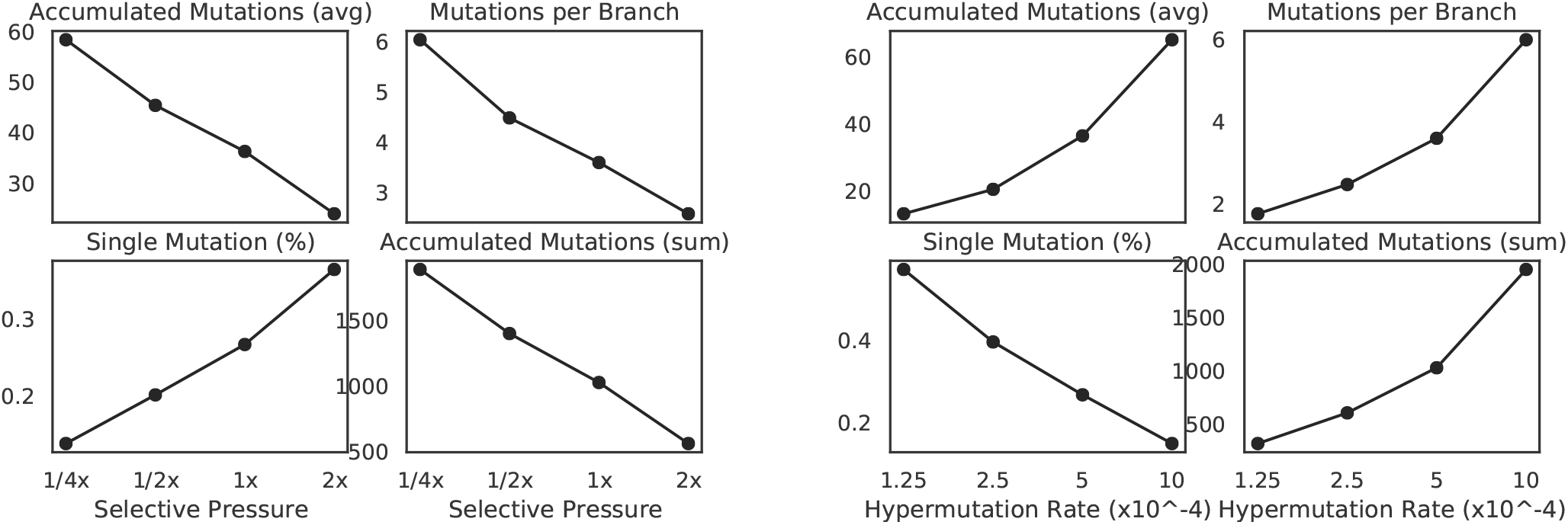
Impact of selective pressure *A* (left) and mutation rate *µ* (right) on sequence-based branch length properties on true trees. *µ* = 5 *×* 10^−4^ in (a-d) and *A* = 0.1 in (e-h).

**Fig. S5.**
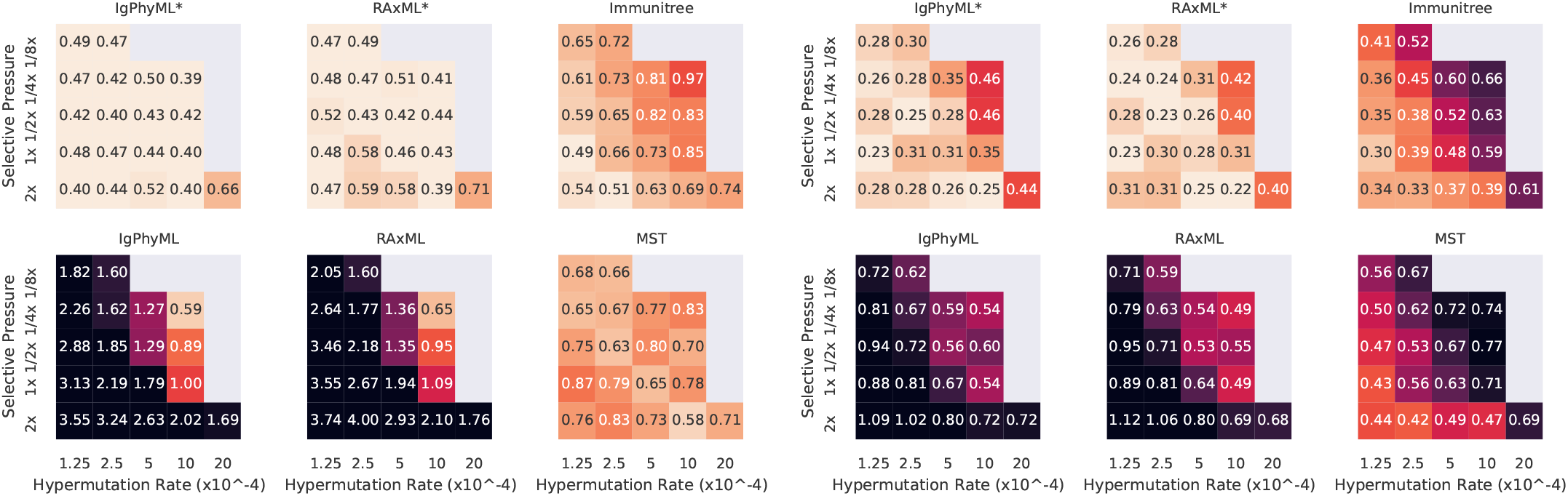
For varying levels of selective pressure (*A*), rate of hypermutation (*µ*), and reconstruction methods, we show MD error (left), and RF error (right). Under some conditions, reconstructed trees from phylogenetic methods are worse than random permuting labels of true tree because both MD and RF (to a lesser degree) severely penalizes resolution of multifurcated nodes.

**Fig. S6.**
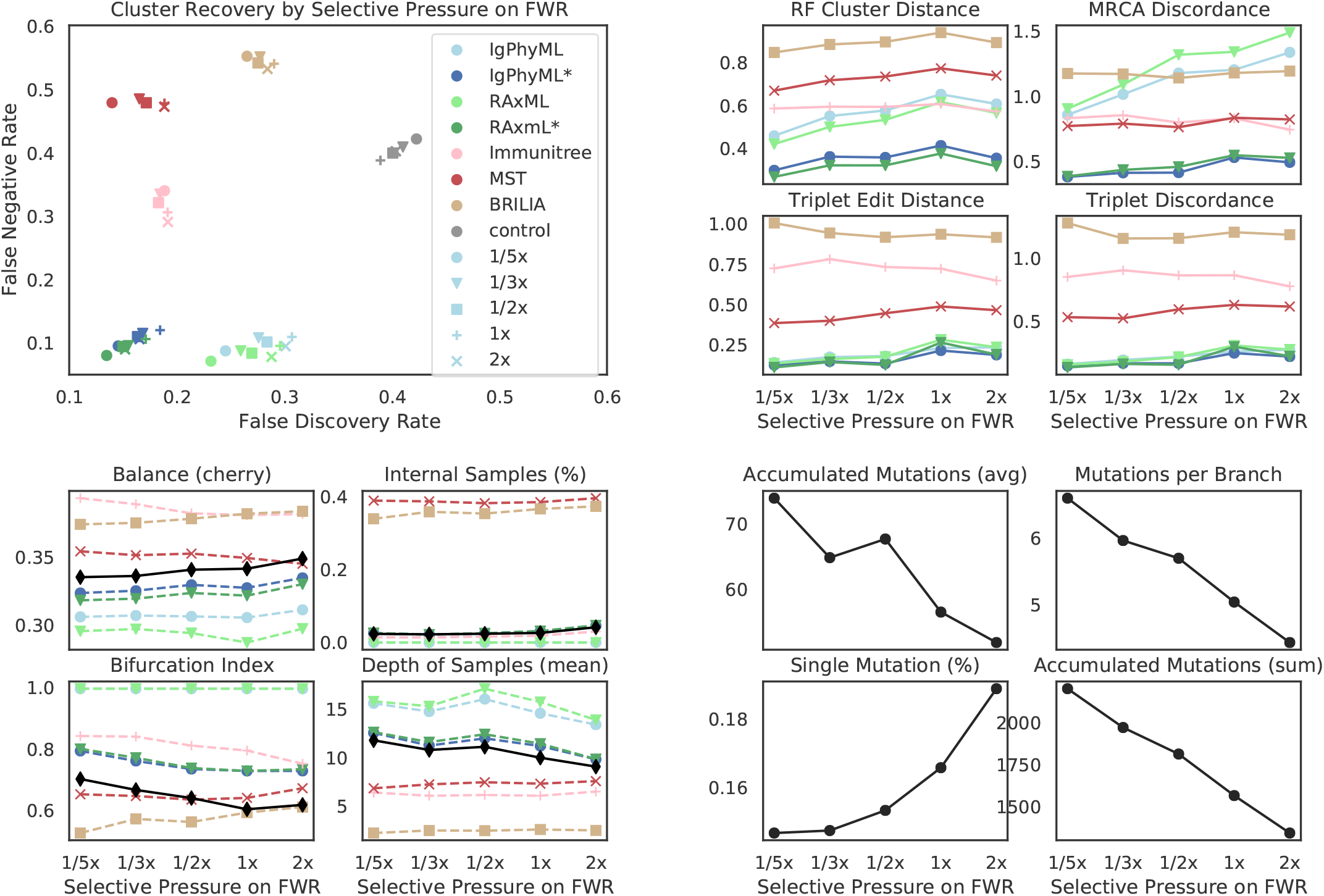
a) FNR versus FDR, b) Robinson-Foulds cluster distance (RF), MRCA Discordance (MD), triplet edit distance (TED), and triplet discordance (TD) by BLOSUM weight multiplier of framework region (*w*_*f*_) and reconstruction methods. c) Properties of true (black) and reconstructed trees by BLOSUM weight multiplier of framework region (FR). d) Properties of true trees.

**Fig. S7.**
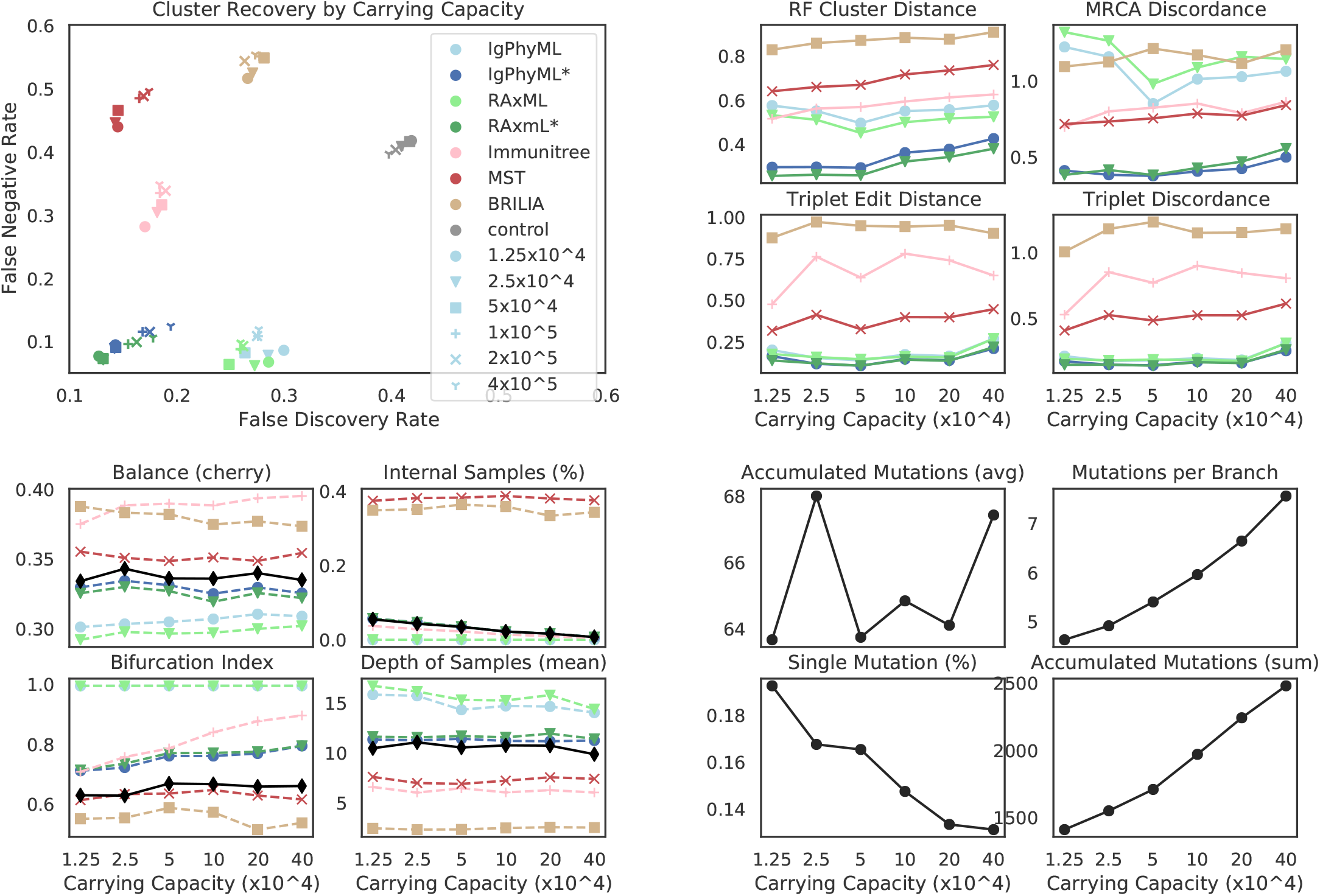
a) FNR versus FDR, b) Robinson-Foulds cluster distance (RF), MRCA Discordance (MD), triplet edit distance (TED), and triplet discordance (TD) by germinal center capacity (*C*) and reconstruction methods. c) Properties of true (black) and reconstructed trees by carrying capacity of germinal center of FR. d) Properties of true trees.

**Fig. S8.**
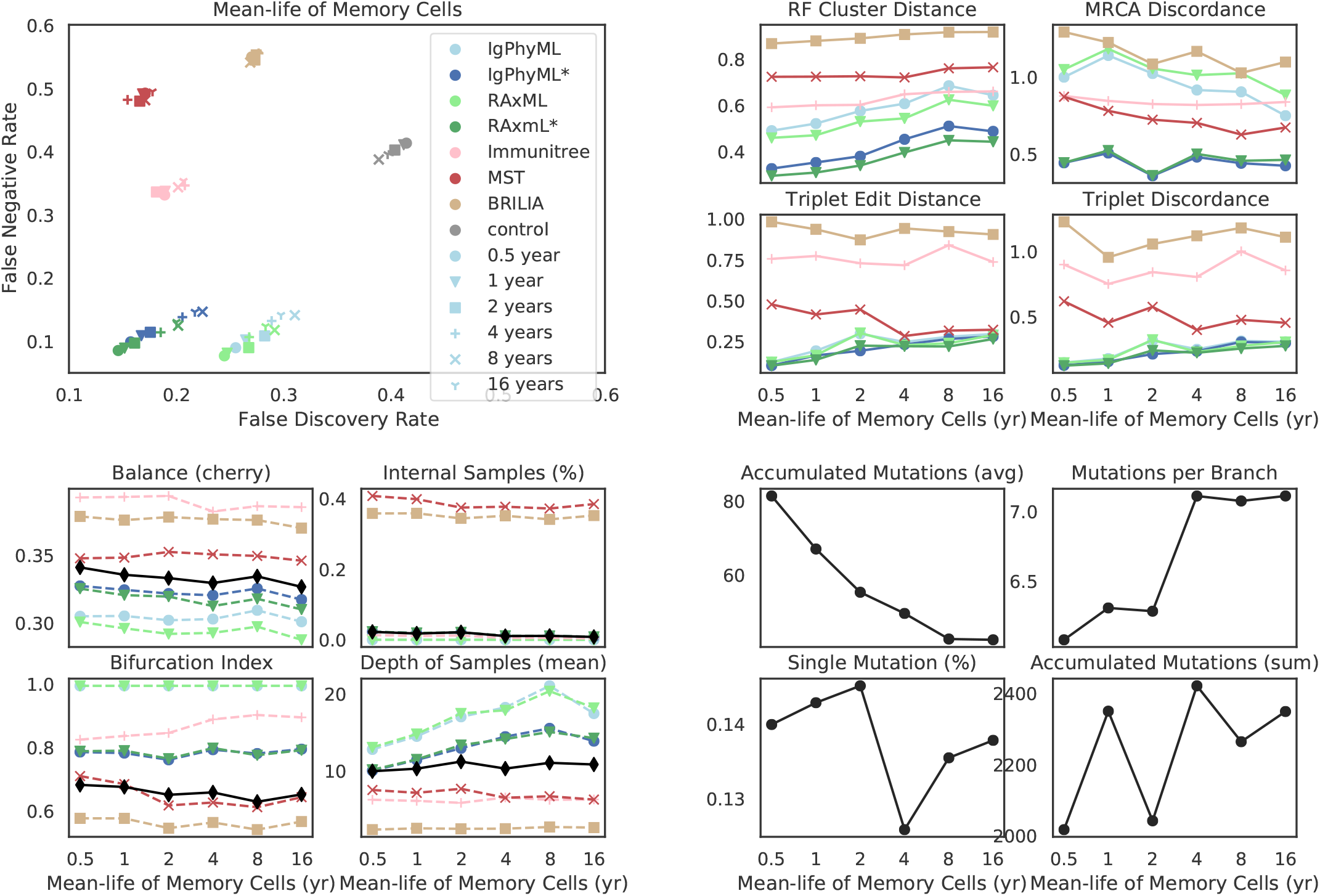
a) FNR versus FDR, b) Robinson-Foulds cluster distance (RF), MRCA Discordance (MD), triplet edit distance (TED), and triplet discordance (TD) by mean memory cell life-time 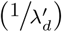 and reconstruction methods. c) Properties of true (black) and reconstructed trees by memory cell life (mean). d) Properties of true trees.

**Fig. S9.**
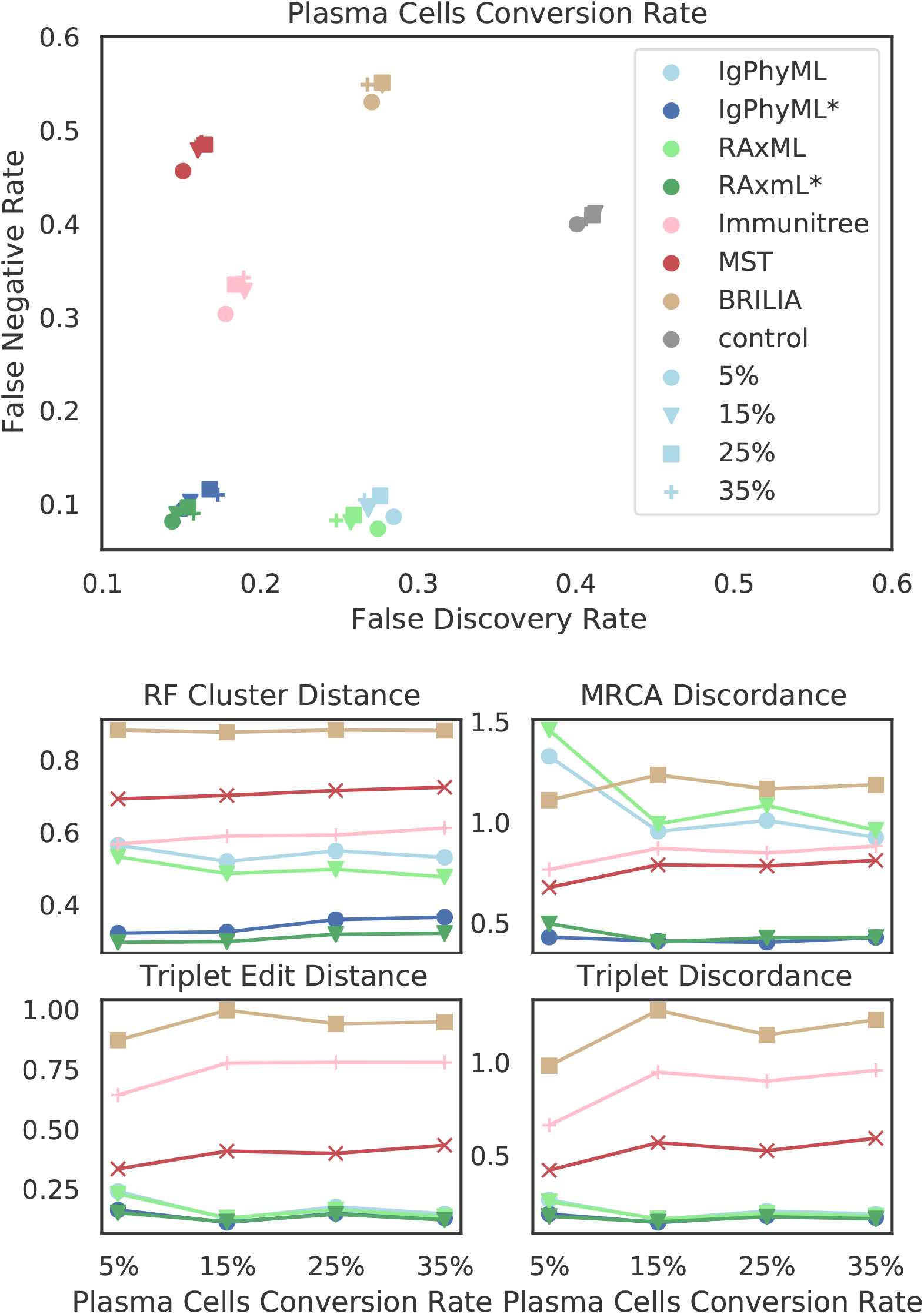
a) a) FNR versus FDR, b) Robinson-Foulds cluster distance (RF), MRCA Discordance (MD), triplet edit distance (TED), and triplet discordance (TD) by fraction of activated cells turning into plasma cell per cell division (*ρ*_*p*_).

**Fig. S10.**
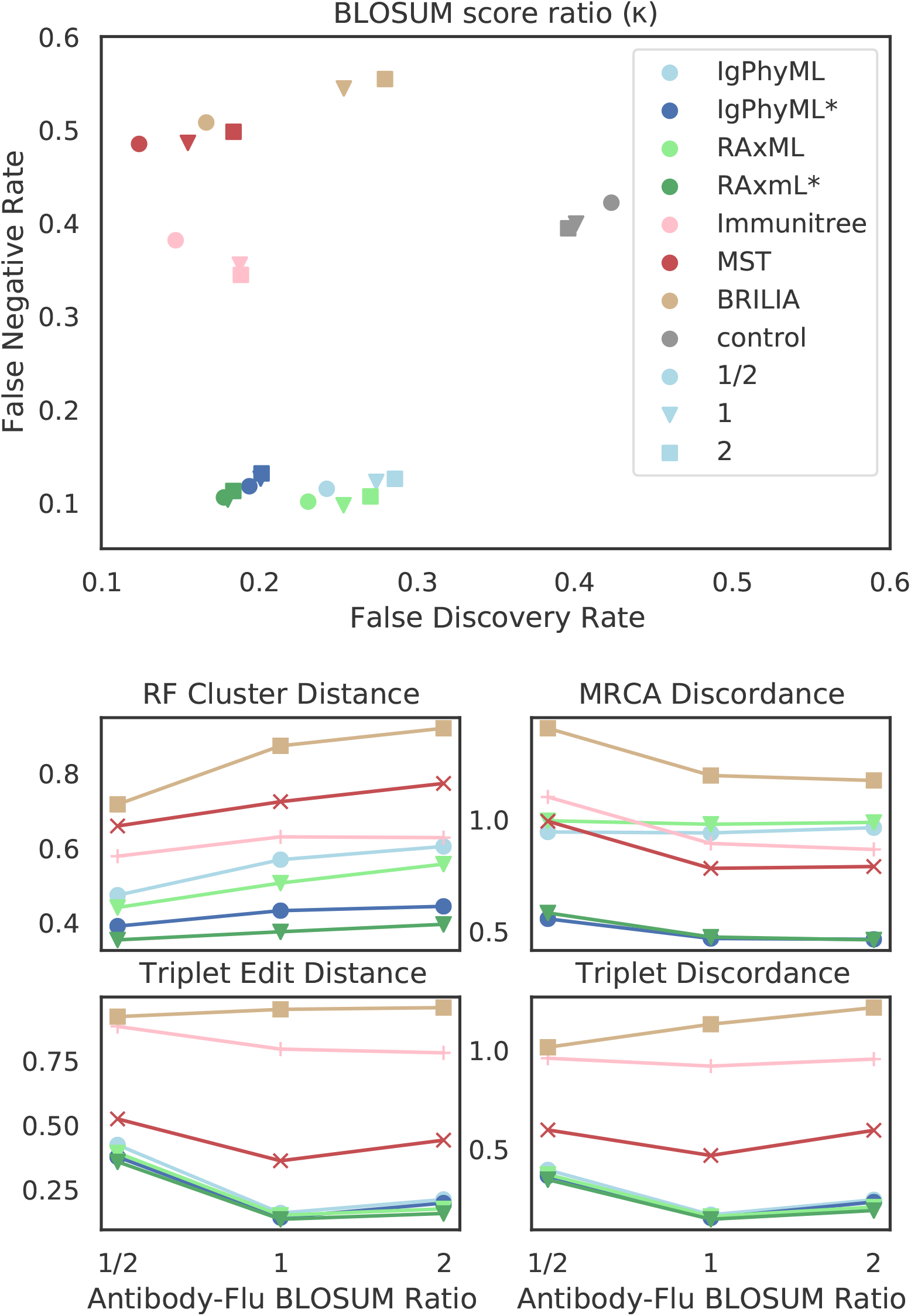
a) a) FNR versus FDR, b) Robinson-Foulds cluster distance (RF), MRCA Discordance (MD), triplet edit distance (TED), and triplet discordance (TD) by BLOSUM score ratio of antibody-coding sequences to antigen sequences (*κ*)

**Fig. S11.**
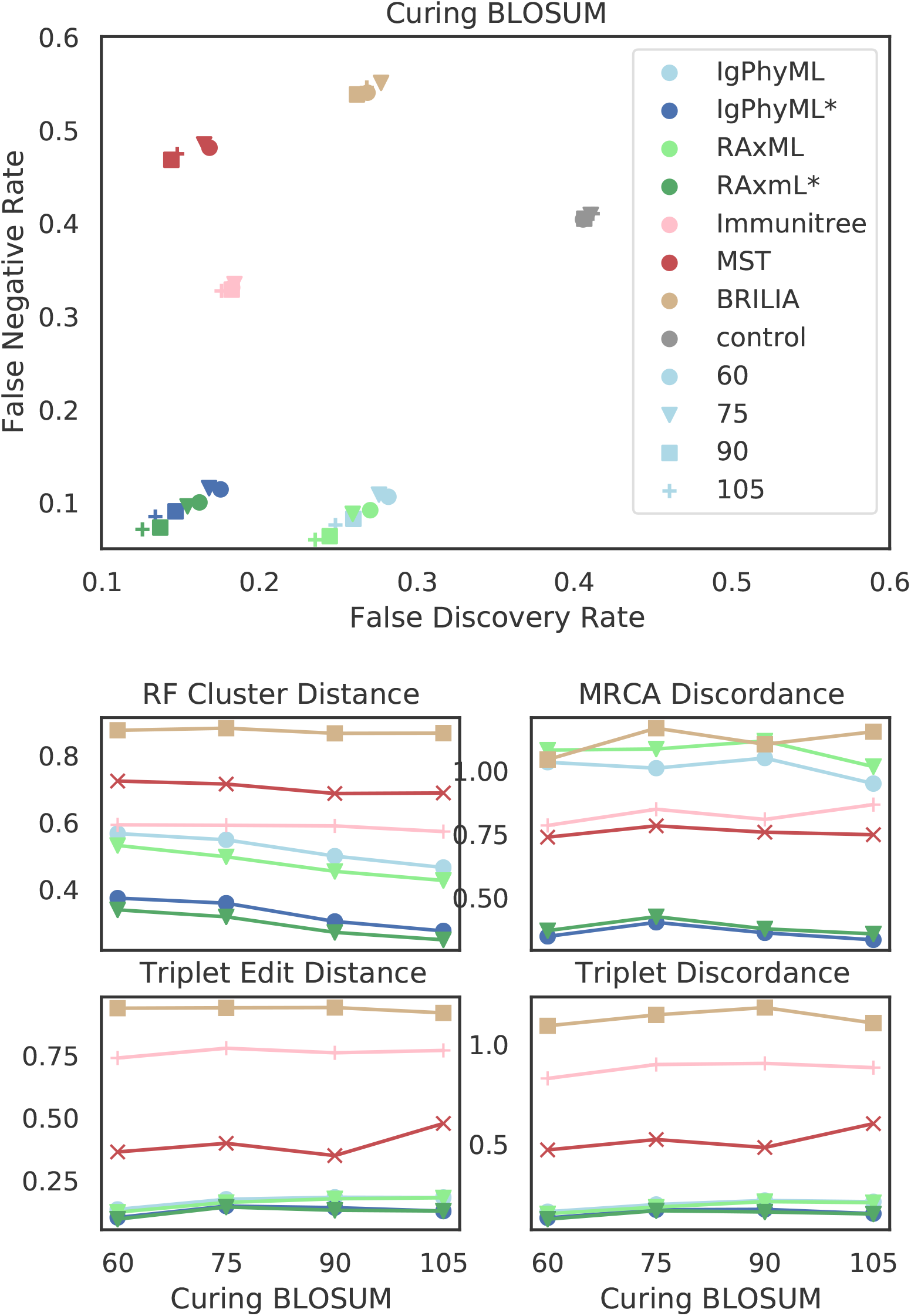
a) a) FNR versus FDR, b) Robinson-Foulds cluster distance (RF), MRCA Discordance (MD), triplet edit distance (TED), and triplet discordance (TD) by BLOSUM score of activated cell antibody-coding sequences that leads to cure 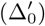.

**Fig. S12.**
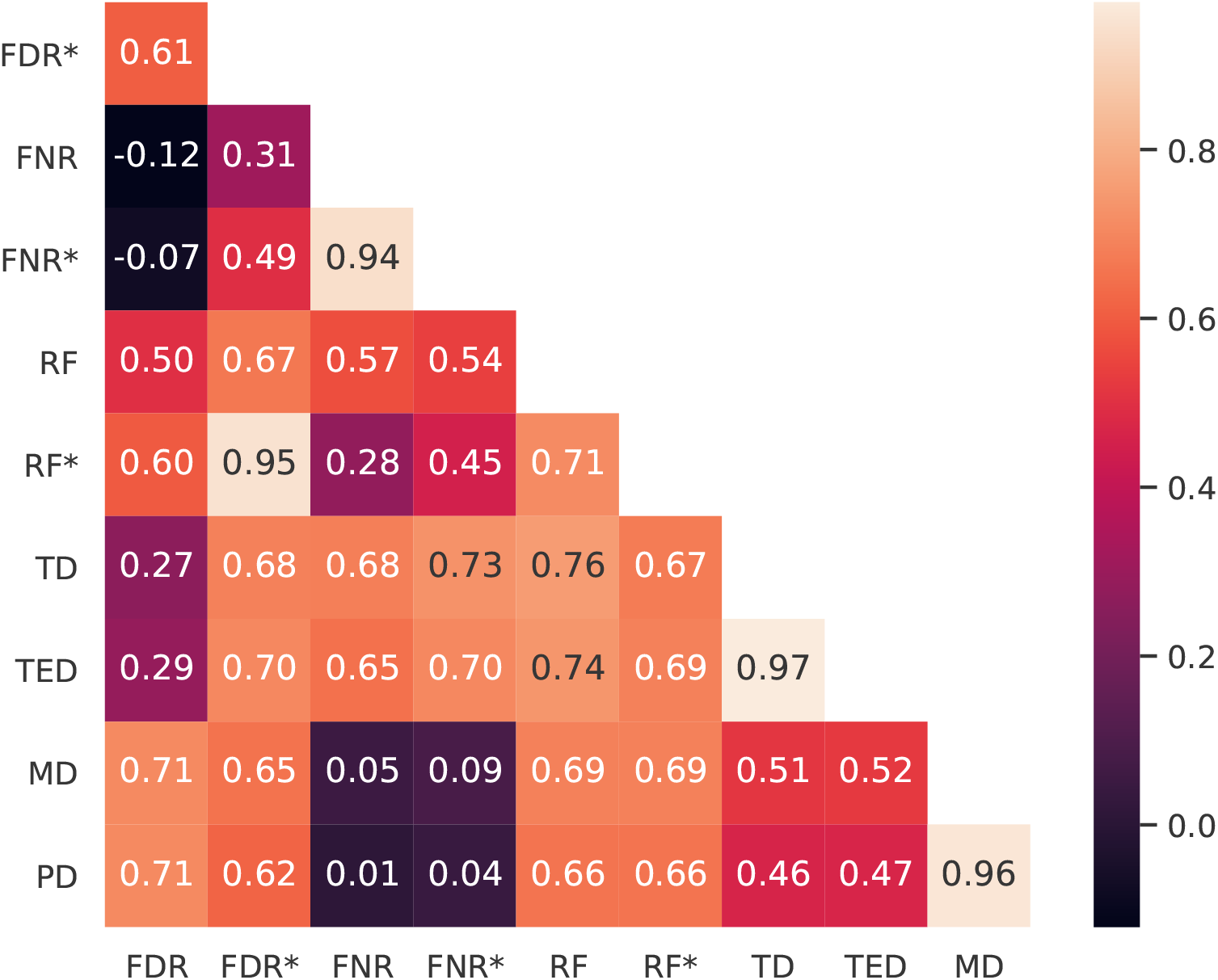
Correlations of evaluation metrics. For each replicate of each simulation condition, we compute Spearman’s rank correlation coefficient of reconstruction method for each pair of evaluation metrics. Here, we show the average coefficient over all replicates of all simulation conditions.

#### Supplementary Algorithms

##### Algorithm S1

Simulating the next event and update time and *S* accordingly. Before running this procedure, we have computed **S** and 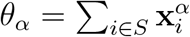 for all *α* from the previous calls to this function (i.e., previous time steps). For each *α*, we have also built an interval tree *T*_*α*_ on leafset *S* and each node *v* storing the summation of 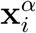 for each leaf *i* under *v*.

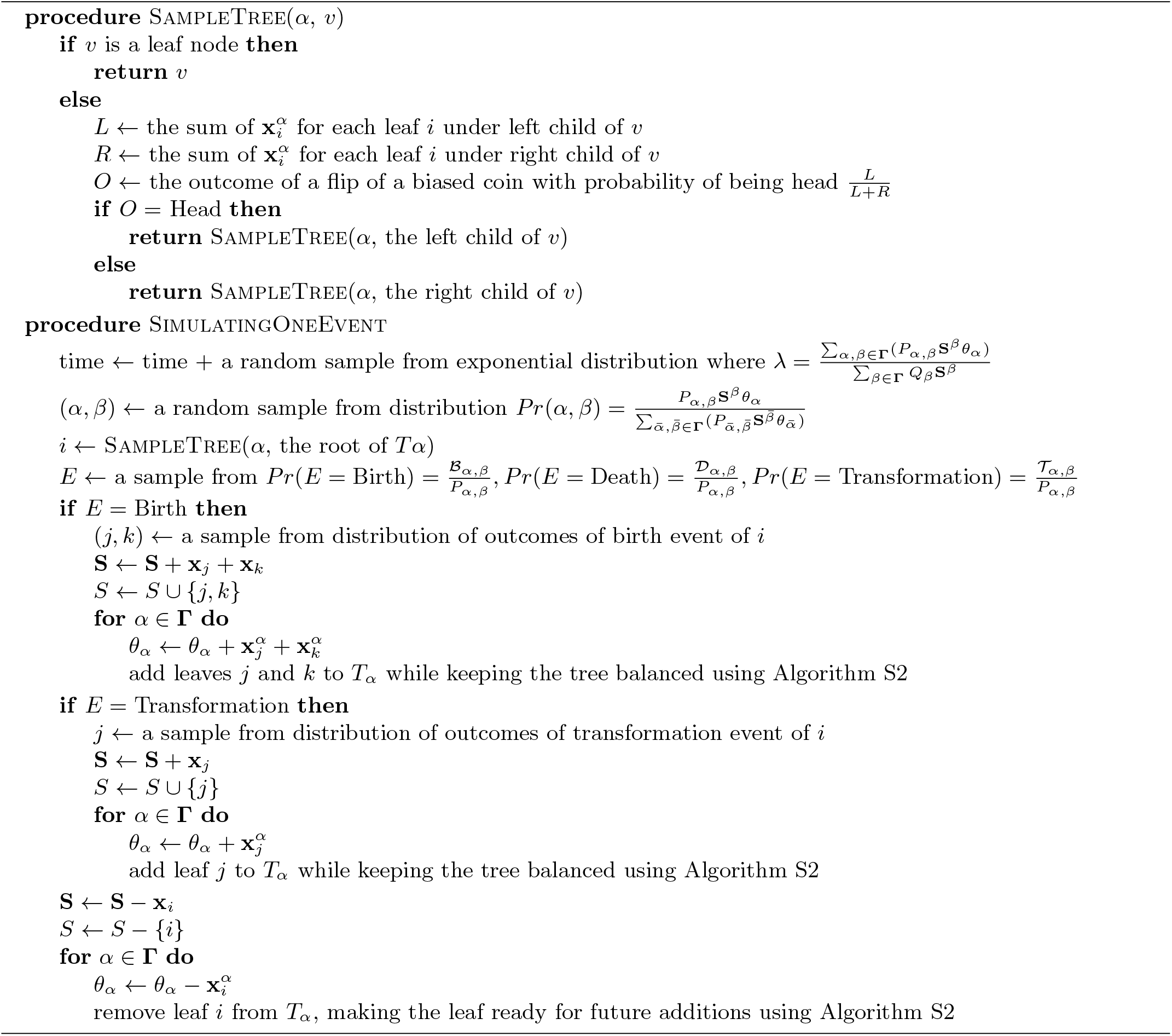

##### Algorithm S2 Exact algorithm for inserting or removing a leaf from tree *T*_*α*_ keeping it balanced. *T*_*α*_ is represented by a full binary tree where each leaf is labeled with either one particle or ∅ and each node *v* has weight *w*_*v*_ equal to the sum of 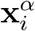 for all leaves under *v* with label (*i*) not being ∅. Assuming a stack *S*_*α*_ keeps all leaves with label ∅.

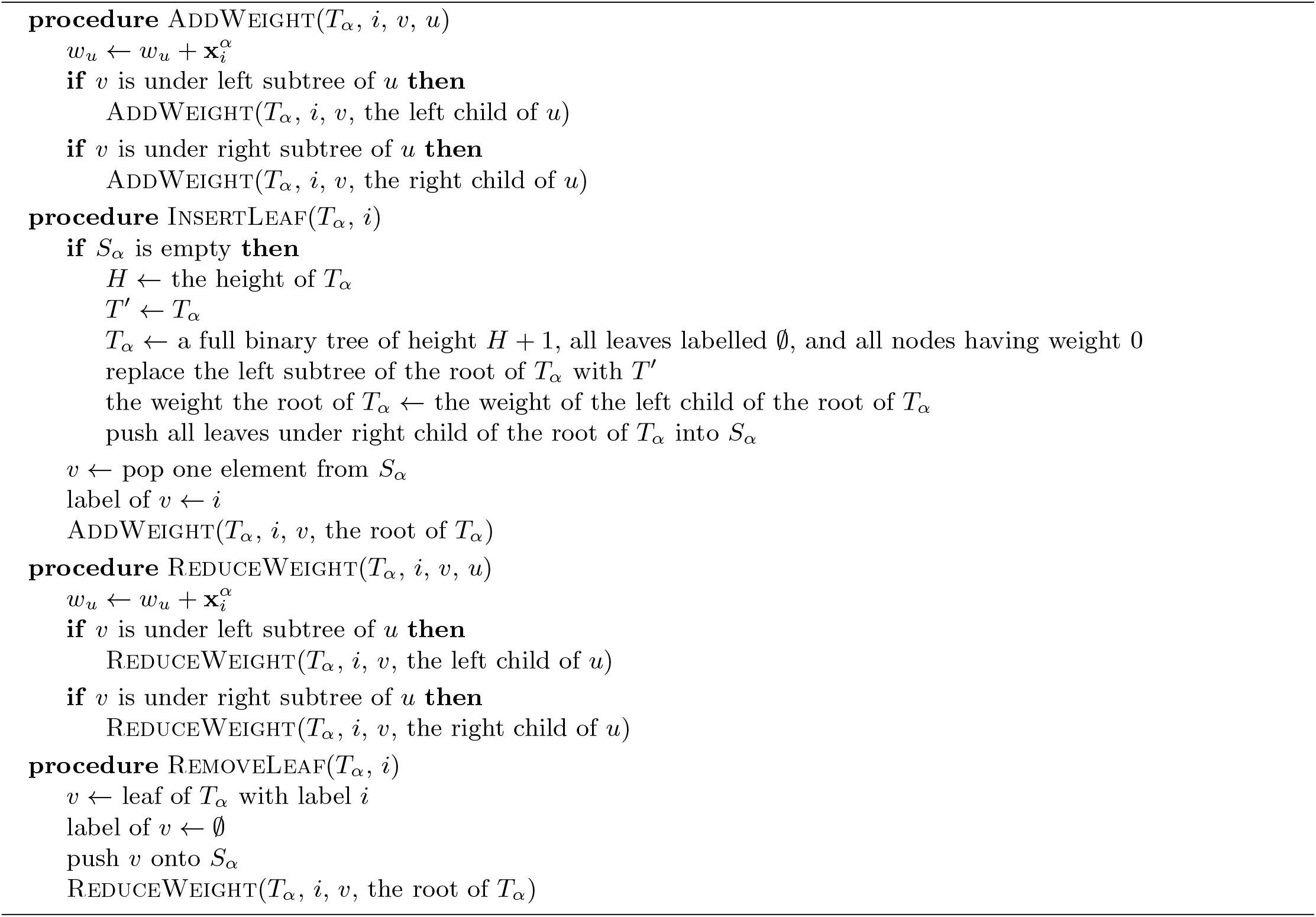

##### Algorithm S3 Heuristics for choosing target sequences to minimize the objective function (3).

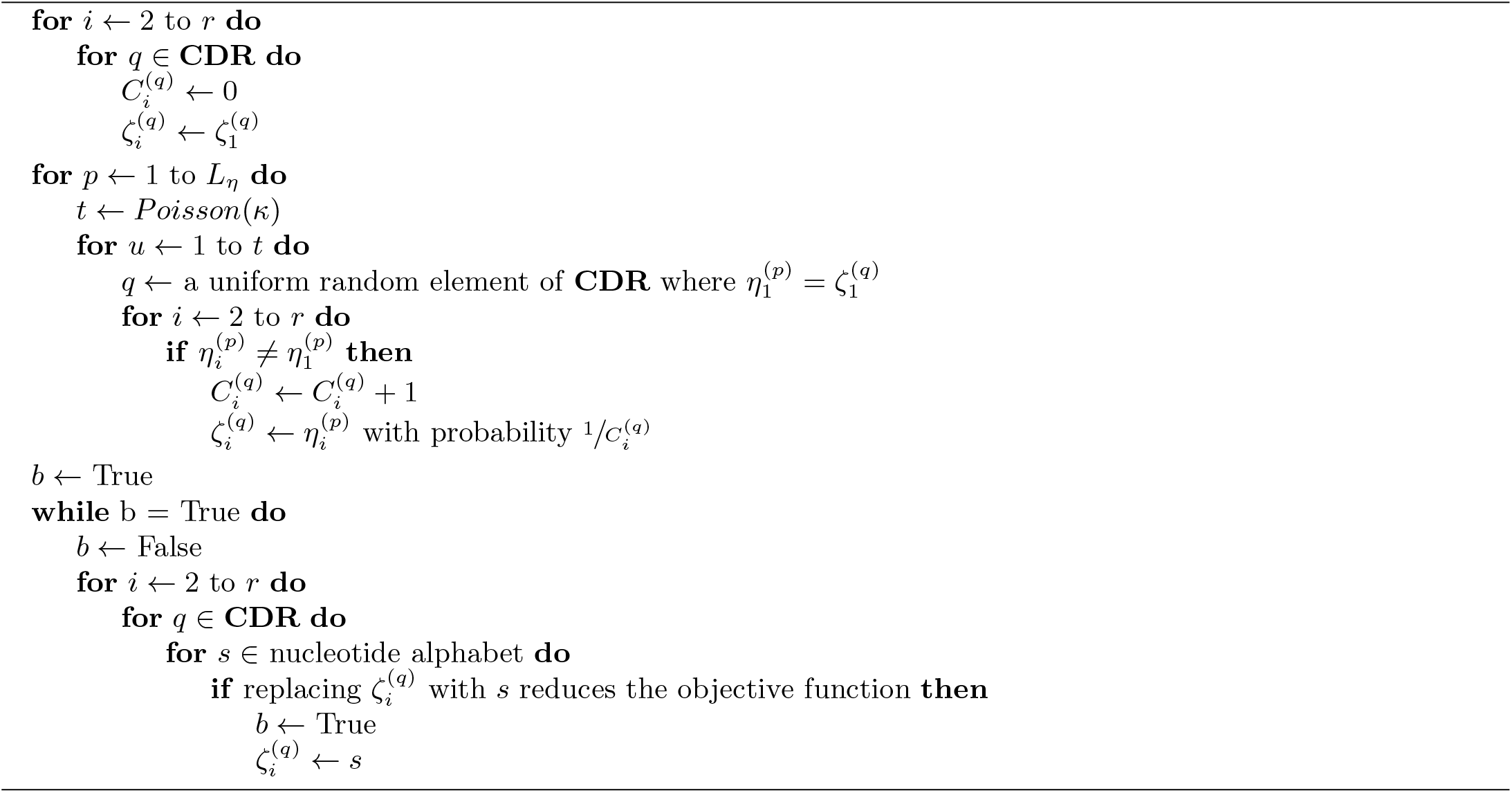

##### Algorithm S4 Let each label be uniformly randomly assigned an element in a finite Abelian group with large enough order (e.g., 64-bit integers). To compute FNR, FDR, and RF, we just need to compute | *ϕ* (*R*) | = |*S*_*R*_|, | *ϕ* (*E*) | = |*S*_*E*_|, and | *ϕ* (*R*) ⋂ *ϕ* (*E*) | = |*S*_*R*_ ⋂ *S*_*E*_|, where set S_T_ for tree *T* can be computed by calling ComputeSet(*T*, the root of *T*).

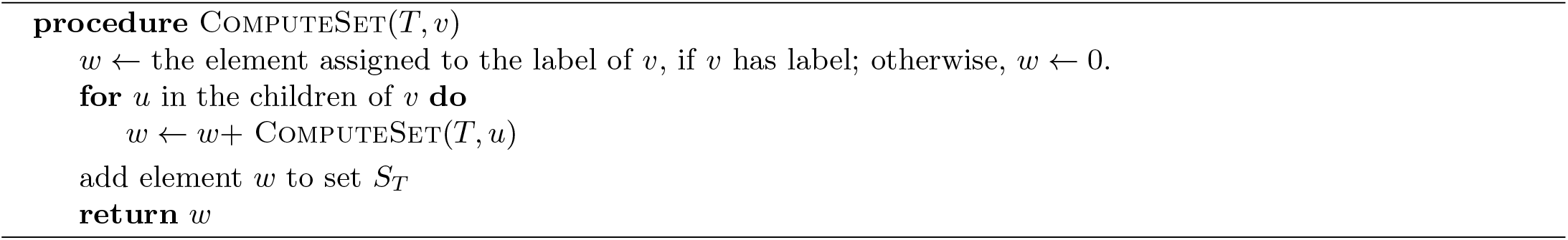

